# Stimulus-Response signaling dynamics characterize macrophage polarization states

**DOI:** 10.1101/2022.03.27.485991

**Authors:** Apeksha Singh, Supriya Sen, Adewunmi Adelaja, Alexander Hoffmann

## Abstract

Macrophages show remarkable functional pleiotropy that is dependent on microenvironmental context. Prior studies have characterized how polarizing cytokines alter the transcriptomic and epigenetic landscape. Here we characterized the immune-threat appropriate responses of polarized macrophages by measuring the single-cell signaling dynamics of transcription factor NFκB. Leveraging a fluorescent protein reporter mouse, primary macrophages were polarized into 6 states and stimulated with 8 different stimuli resulting in a vast dataset. Linear Discriminant Analysis revealed how NFκB signaling codons compose the immune threat level of stimuli, placing polarization states along a linear continuum between the M1/M2 dichotomy. Machine learning classification revealed losses of stimulus distinguishability with polarization, which reflect a switch from sentinel to more canalized effector functions. However, the stimulus-response dynamics and discrimination patterns did not fit the M1/M2 continuum. Instead, our analysis suggests macrophage functional niches within a multi-dimensional polarization landscape.

**Highlights:**

- Polarization of macrophages affects stimulus-response NFκB dynamics
- For each condition, NFκB signaling codons quantify the “immune threat” level
- Machine Learning reveals polarization-induced canalization of stimulus-responses
- NFκB stimulus-responses may define a landscape of macrophage states

**eTOC blurb:** Macrophages are profoundly responsive to their tissue microenvironment, but how that affects their pathogen response functions has not been investigated systematically. Here we studied how their signaling response is affected by six polarizing cytokines. We found each modulates their stimulus-responses highly specifically, producing distinct patterns of stimulus-discrimination. Thereby, these stimulus-response specificities may be used to describe a landscape of functional macrophage states.

**Graphical Abstract:** 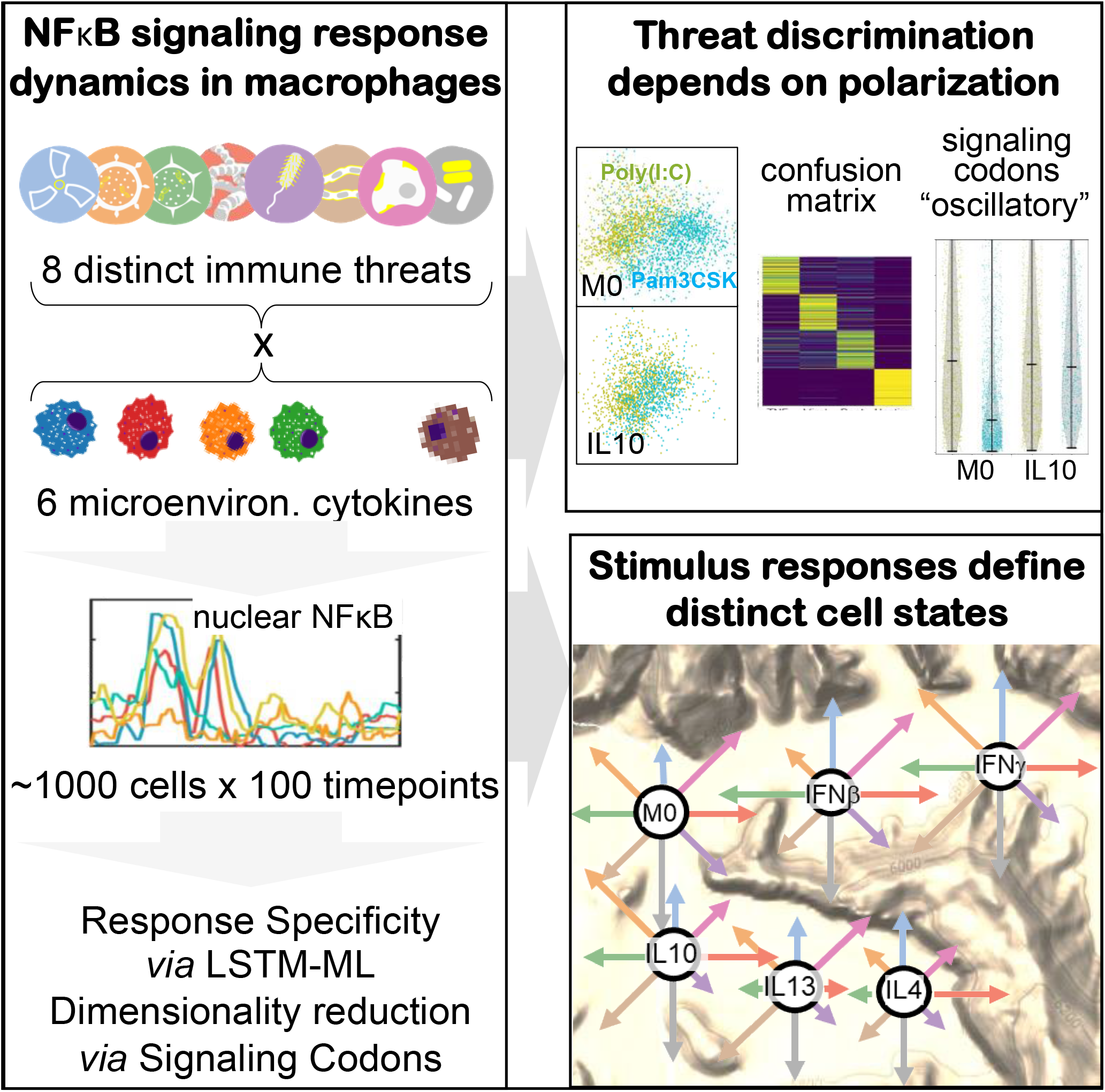

## INTRODUCTION

Macrophages have critical functions in the immune response. Upon detecting pathogen or tissue damage through pattern recognition receptors or cytokines through cognate receptors, these cells can perform a wide range of tasks from the phagocytosis of pathogen components and cellular debris to antigen presentation, recruitment of other immune cells to sites of infection, and activation of system-wide immune responses. Which functional responses are elicited depends not only on the activating stimulus, but also on the microenvironmental context of the macrophage^1^. More specifically, macrophages are polarized into different biological functional states by the microenvironmental cytokine milieu to accentuate specific functional responses over others^2^.

Macrophage polarization was first described in terms of a M1 vs. M2 dichotomy, although it is now recognized that these states are representative of a larger spectrum of macrophage activation *in vivo*^3^. M1 macrophages found in inflamed microenvironments defined by the presence of IFNγ, play critical roles in defending the host from pathogens, such as in bacterial, viral, and fungal infections. Alternatively activated or M2 macrophages have anti-inflammatory function and additionally regulate wound healing and repair functions^4^. Subsets of M2 macrophages have been described, such as M2a promoted by IL13 and IL4 exposure and M2c promoted by IL10 exposure^5^. Abnormalities in macrophage activation and subsets of polarized cells have been implicated in disease such as metabolic disorders, asthma, allergic reactions, cancer, and autoimmune disorders^1,6,7^. Many previous studies have attempted to characterize differences in polarization states based on transcriptomic^8–10^, epigenomic^11,12^, or proteomic^13,14^ profiling, with recent advances in single-cell technologies revealing heterogeneity within these states^15–17^. However, functional states of macrophages ought to be defined by their actual functionality. Steady-state measurements of molecular abundances provide correlative markers of these states, but profiling single-cell functional responses may reveal a state map that is closer to their biologically relevant functions

Macrophages must recognize and react to diverse stimuli to fulfill their biological roles. Macrophages not only need to detect different pathogen or host stimuli, but need to mount a response that is appropriate to the stimulus encountered^17–19^. The signaling system that controls macrophage responses to pathogen, tissue injury, or cytokine activates a handful of effectors, including the central immune response transcription factor, NFκB. NFκB activation shows stimulus-specific activation dynamics^20–24^ that can control the expression of immune response genes^25–28^ and reprogram the epigenome^29^. A recent set of single-cell studies in primary macrophages has characterized a temporal signaling code that consists of six dynamical features, termed “signaling codons”, that are deployed stimulus-specifically^30^.

Indeed, this “NFκB response specificity” may be quantified with information theoretic or machine learning classification approaches. Diminished response specificity was found to be associated with macrophages from a mouse model of the autoimmune disease Sjögren’s syndrome.

However, whether and how the stimulus-specificity of signaling codon deployment is affected by polarizing cytokines remains unexplored, as well as the potential of using NFκB response dynamics to map macrophage polarization states.

Many studies have described molecular mechanisms by which polarizing cytokines affect NFκB activation, but what their consequence is for macrophage response specificity in different contexts remains unclear. Type 1 interferons, such as IFNβ, inhibit the translation and promote degradation of IκBα and increase expression of receptors like RIG-1 which activate IKK^31,32^. Type 2 interferons, such as IFNγ, induce PA28 proteins which enhances the degradation of free IκBα^32^ or IκBε^33^. In this way, IFNβ and IFNγ may alter NFκB activation dynamics. IL4 and IL13 stimulation results in STAT6 activation and downstream KLF4 expression that sequesters coactivators required for NFκB activation^6^. IL10 stimulation induces p50 NFκB homodimers^34^. Multiple microRNAs (miRNAs) have been identified as regulators of TLR signaling components and whose expression can be modulated by polarizing cytokines as well^35,36^. IL10 is necessary for the expression of miRNA-146b which negatively regulates TLR4 signaling^37^. M1 polarization increases the expression of miRNA-155, which targets MYD88^38^, whereas M2 polarizers decrease its expression^39,40^. Finally, the expression of miRNA-146a, which targets TRAF6, is responsive to many inflammatory stimuli, including interferons, while IL4 reduces its expression^41,42^. While there is a rich literature of molecular mechanisms engaged by polarizing cytokines, it remains unclear how polarization affects the biologically relevant properties of the transcriptional effectors of the macrophage signaling system, such as their stimulus-specific dynamics and the resulting response specificities.

Here, we examined how macrophage polarization affects stimulus-specific dynamics of NFκB activation by leveraging a live microscopy workflow to generate a large dataset of single-cell nuclear NFκB timecourse trajectories in response to 8 stimuli and 6 polarization conditions. We applied machine learning approaches to decompose NFκB responses and quantitatively characterize NFκB response specificity across polarization states. This analysis revealed that NFκB signaling codons define the immune threat level of a response and that this level is a function of the macrophage polarization state. Furthermore, utilizing a Long Short-Term Memory (LSTM) based machine-learning classifier, we identified stimuli responses that were less distinct with polarization, such as host TNF vs. pathogen ligands, and viral vs. bacterial ligands. Such convergence is associated with changes in signaling codon deployment that shift the immune threat level. Finally, we used the rich dataset of stimulus-specific NFκB response dynamics to generate multi-dimensional mappings of macrophage polarization states. The polarization-specific differences in NFκB dynamics and resulting differences in response specificity suggest a specialization of macrophages into distinct functional niches.

## RESULTS

### An experimental pipeline for studying stimulus-specificity in polarized macrophages

To study how polarization of macrophages by microenvironmental cytokines may affect NFκB signaling responses to various pro-inflammatory stimuli, we sought to generate a large dataset with mVenus-RelA knockin macrophages that were polarized in 6 different conditions, and then stimulated with 8 different proinflammatory stimulation ligands. Generating this large dataset with 48 experimental conditions was made feasible by producing macrophages from a HoxB4-immortalized myeloid precursor line^43^ derived from the mVenus-RelA knockin mouse strain.

Macrophages produced in this manner showed responses that were close to indistinguishable from those observed in bone marrow-derived macrophages in terms of NFκB signaling dynamics and endotoxin-induced gene expression in contrast to the often-used Raw264.7 cell line (Figure S1).

Within our experimental workflow, differentiated macrophages were exposed to either IFNβ or IFNγ to polarize towards M1, or IL10, IL13, or IL4 for M2 polarization, and then stimulated with agonists for different toll-like receptors such as R848 (TLR8), Poly(I:C) (TLR3), Pam3CSK (TLR1/2), CpG (TLR9), Flagellin (TLR5), FSL1 (TLR2/6), or LPS (TLR4) as well as the pro-inflammatory cytokine TNF (Figure 1A). The polarizing conditions did not appear to dramatically impact the macrophages’ morphology visually (Figure S2). The resulting single-cell nuclear NFκB trajectories were captured by an established live-cell microscopy workflow and quantified by a robust image analysis pipeline ^30^. For each experimental condition, we obtained two biological replicates, with hundreds of single-cell NFκB trajectories that passed stringent quality control metrics (see Methods) in each dataset (Figure 1B). This dataset encompasses a total of 68,056 cells, each characterized by 98 microscopy images.

**Figure 1:**
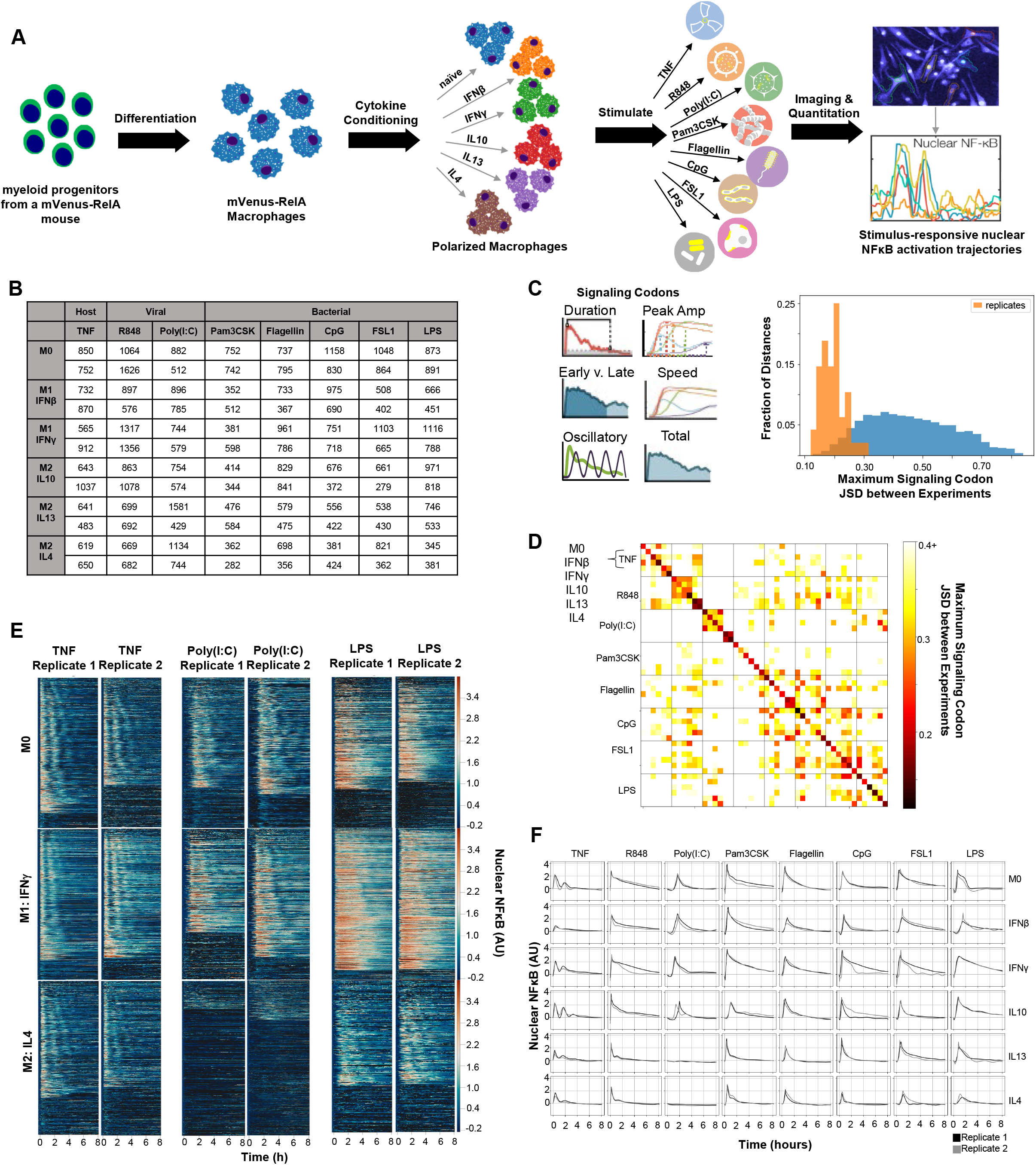
Single-cell NFκB trajectories across 6 polarization states following 8 different stimulations. **A**. Experimental pipeline for obtaining single-cell NFκB responses in different polarization and stimulation conditions to study the effect of polarization on stimulus response specificity **B**. Number of single cell NFκB trajectories obtained in each experimental condition with two biological replicates **C**. Histogram of maximum Jensen-Shannon Distance (JSD) between distributions of Signaling Codon (SC) from experiments, with distances between replicate experiments in orange and distances between different experimental conditions in blue. **C**. Along the diagonal of the distance matrix are the maximum SC JSD between replicates for each experimental condition. Experimental conditions are ordered by stimuli and further sub-ordered by polarization state. The off-diagonal elements are the mean maximum SC JSD between replicates of different experimental conditions. **E**. Example replicate NFκB trajectory datasets in M0, M1:IFNγ, and M2:IL4 polarization states with TNF, Poly(I:C), and LPS stimulation. Each row in a heatmap corresponds to a single macrophage in the experiment and the color corresponds to the amount of nuclear (active) NFκB. **F**. Soft-DTW (dynamic time warping) barycenter of all NFκB trajectories in each replicate for all experimental conditions (computed using smoothing hyperparameter γ = 5).

We examined the replicates by focusing on previously identified trajectory features, termed signaling codons (Figure 1C). Using the Jensen-Shannon distance (JSD) of signaling codons between each population of cells as a measure of similarity, we found that the maximum JSD between replicates were in general much smaller than between cells stimulated in different conditions. This assures that the biological differences of interest are larger than the technical variability associated with the experimental and image analysis workflow. A more detailed analysis revealed that some polarization and stimulus combinations to be more similar than most (Figure 1D), such as responses to R848 in cells polarized with IFNγ or IL10, or responses to TNF in cells polarized with IFNγ and responses to R848 when polarized with IL13 or IL4.

Visual inspection of heatmaps that depict the actual time-course measurements confirmed that stimulus-specific signaling characteristics are preserved in each replicate, while the precise fraction of non-responding cells varied between some replicates (Figure 1E).

To visualize the trajectories in an aggregate form for each condition, the soft-DTW (Dynamic Time Warping) barycenter^44,45^ of the NFκB trajectories in each replicate was computed^46^ (Figure 1F). A barycenter is a constructed trajectory that minimizes the pairwise distance between itself and each trajectory in the input dataset. Even in aggregate form, NFκB dynamics showed stimulus-specificity, most notably for TNF, as well as a degree of polarization specificity, such as a loss in response to Poly(I:C) with IL13 and IL4 polarization. While this analysis confirmed the reproducibility of replicates in visual form, it is also apparent that the full dynamic features observed in single-cell trajectories are lost in the aggregation.

With the quality of this rich dataset of single-cell NFκB trajectories confirmed, we turned to computational data analysis methods that respect the single-cell nature of the data to characterize the effect of polarization on the macrophage’s stimulus-responses.

### The immune threat level of a stimulus is modulated by polarization

Prior work suggested that the level of immune threat a macrophage encounters is encoded by responsive NFκB signaling dynamics^47^. Here we explored whether the immune threat level encoded by the macrophage to a particular stimulus is a function of the polarization state. We sought to establish how the aforementioned signaling codons could define the level of immune threat by determining how they compose a host TNF vs. a pathogen response. We used Linear Discriminant Analysis (LDA) to find a linear combination of the signaling codons that attempts to discriminate all TNF from all PAMP responses in our dataset (Figure 2A). Under this LDA projection, pathogen responses were associated with higher positive values, thereby defining the immune threat level of a response (Figure 2B). Utilizing the projection, we found that the mean response to LPS with IFNγ polarization had the greatest immune threat characterization, whereas the mean response to TNF in the naïve condition had the least. This result supports the notion that maximal macrophage activation is elicited by LPS plus IFNγ^21,30,32^ and that TNF secreted by other cells represents a lower immune threat level than direct cellular exposure to PAMPs.

**Figure 2:**
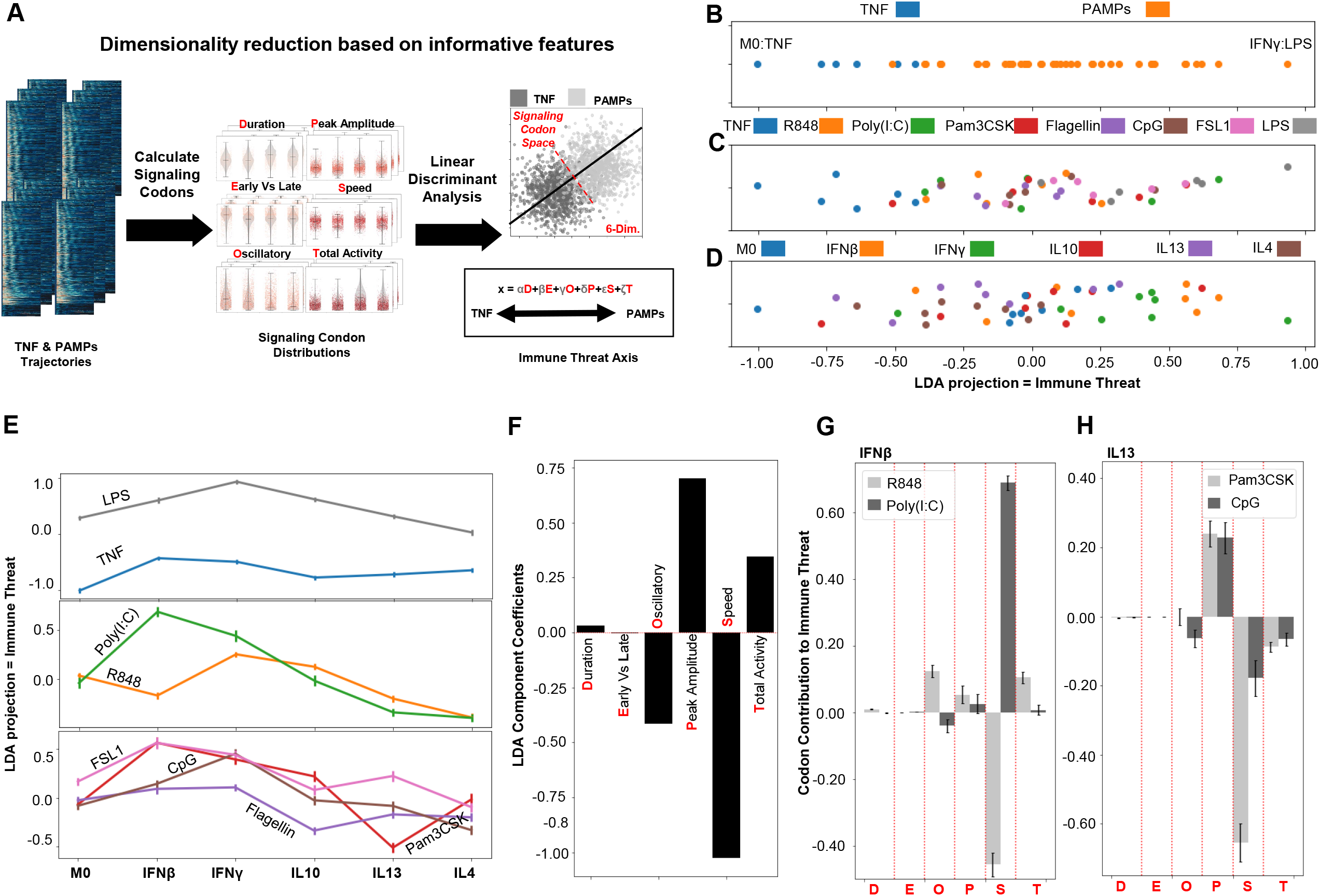
Characterization of Immune Threat utilizing LDA of Signaling Codons. **A**. Signaling Codons are calculated from the single-cell NFκB trajectories and then LDA finds a linear combination that best distinguishes threat level **B**. Comparing mean LDA projection of host TNF versus pathogen (PAMPs) responses; this axis quantifies immune threat as pathogen responses are more positive along it. **C**. Comparing mean LDA projection of different ligand responses **D**. Comparing mean LDA projection of different polarizer responses shows more mean M1 polarized responses with positive LDA values and more mean M2 polarized responses with negative LDA values. **E**. Mean immune threat level of each stimuli versus polarization state shows modulation by polarizers **F**. Coefficient applied to each signaling codon to obtain the LDA projection, hence characterizing immune threat: decreased oscillations, increased peak amplitude, decreased speed, and increased total activity. **G**. Scaled mean signaling codon values for viral ligands, R848 and Poly(I:C), in IFNβ polarization condition, hence specifying codon contribution to immune threat quantification. **H**. Scaled mean signaling codon values for bacterial ligands, Pam3CSK and CpG, in IL13 polarization condition. Error bars in E, G, and H correspond to 95% confidence intervals.

When we examined how the eight macrophage stimuli aligned on the LDA projection, we found indeed that, as expected, the host stimulus TNF consistently scored at the low end, but that the responses to PAMPs were variable (Figure 2C). Mean responses to Poly(I:C), for example, ranged from −0.39±0.02 to +0.68±0.03, and even mean responses to LPS ranged from 0.03±0.03 to 0.93±0.02. This suggests that polarization states modulate the perceived immune threat level encoded in stimulus-specific NFκB responses. Indeed, examining how polarization states align on the LDA projection, we found that responses associated with specific polarizing cytokines were over-represented in specific parts of the projection (Figure 2D). For example, we found that the highest immune threat values were associated with IFNβ and IFNγ polarization states, whereas negative immune threat values often derived from IL13 and IL4 polarization states, with most responses from unpolarized naïve macrophages clustering near zero. Thus, polarization states were a significant determinant of what immune threat-level macrophages perceive with PAMPs. Furthermore, the analysis supports the notion that macrophage polarization states do not merely fall into two discrete classes of M1 and M2, but may be represented on a continuum.

Plotting how polarization affects the macrophage’s perception of specific stimuli (Figure 2E), we found that while TNF showed a consistently low immune threat level, LPS, which was highest with IFNγ, was diminished to average with IL4. Other PAMPs showed an even greater degree of polarization-dependent variability, with polarization affecting the immune threat characterization of each differently. For example, among the two viral PAMPs, the immune threat level of R848 was maximized with IFNγ polarization but that of Poly(I:C) was maximized with IFNβ polarization. There are also differences between bacterial PAMPs, such as Pam3CSK which has minimal immune threat characterization with IL13 polarization, and CpG which instead has minimal immune threat characterization with IL4 polarization.

We wondered which dynamical features of NFκB signaling were driving the differences in immune threat evaluation. Investigating the coefficients applied to the six signaling codons to generate the linear projection identified by LDA, we found that responses with increased immune threat levels are associated with decreased oscillations, increased peak amplitude, decreased speed, and increased total activity (Figure 2F). For each trajectory, the signaling codon values were multiplied by these coefficients and then summed to obtain the immune threat level value. This provides a way of quantifying the contribution of a particular signaling codon to the immune threat of a specific experimental condition. Comparing the viral ligands, R848 and Poly(I:C), we found that the maximization in Poly(I:C)’s immune threat level with IFNβ polarization was driven by speed (Figure 2G). Similarly, the minimization of the immune threat level of bacterial ligand Pam3CSK with IL13, as opposed to CpG with IL13, was also driven by speed (Figure 2H). Overall, this analysis revealed that the immune threat characterization of stimuli is a function of polarization and we identified specific signaling codons that may drive such changes. Further, the fact that each polarizing cytokine had differential effects on different PAMPs suggested that the macrophage polarization is not adequately described as falling on a linear continuum between M1 and M2 states but there is a more complex multi-dimensional landscape of polarization states.

### A machine learning classifier reveals reductions in stimulus-response specificity with polarization

Given the plasticity of the immune threat level associated with each stimulus, we asked whether polarization may then affect the degree of stimulus-response specificity in NFκB dynamics. To quantify stimulus distinguishability based on NFκB trajectories and characterize how it is affected by polarization, we implemented a Long Short-Term Memory (LSTM)-based machine learning classifier^48^. LSTM is a recurrent neural network (RNN) architecture developed to handle the vanishing/exploding gradient problem frequently encountered when training RNN’s. LSTM networks are well suited to perform classification or prediction tasks on time-series data because of their ability to learn long-term dependencies in input sequences^49^.

The classifier was trained on different ligand identification tasks using 80% of the stimulus-responsive trajectories from all polarization states as input data (Figure 3A, see Methods). By comparing the output model’s classification performance on the remaining 20% of the data, which was unseen during training, we were able to quantify how stimulus distinguishability was affected by polarization (Figure 3A). For each classification task, the data was resampled and the training procedure was repeated 15 times to estimate uncertainty in the obtained performance metrics. To quantify classification performance, two metrics were used (Figure 3B). First, the F1 score is the harmonic mean of the accuracy and precision for each class; it is a measure of classification performance, and hence stimulus distinguishability. Second, the confusion fraction is the mean incorrect prediction probability between pairs of classes and thus quantifies the convergence of the NFκB trajectories associated with two stimuli. We observed that the LSTM-based classifier achieved better performance than an ensemble of decision trees algorithm using the time-series data (Figure S3A).

**Figure 3:**
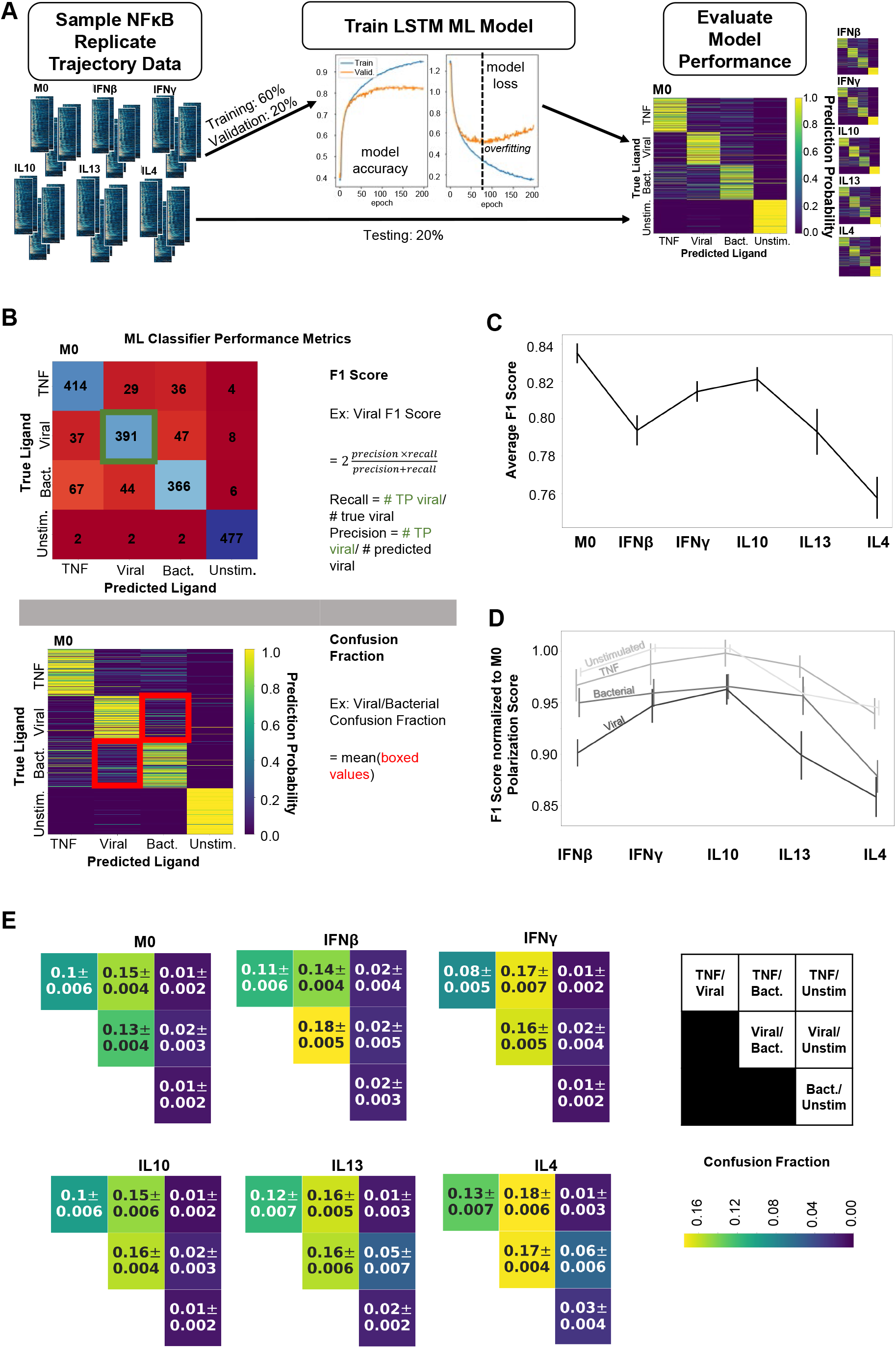
An LSTM-based ML classifier reveals the effect of macrophage polarization on stimulus response specificity. **A**. For each classification task, data was sampled from all polarization states to train and test the LSTM-based ML model. Input data was split into training (60%), validation (20%), and testing sets (20%) where validation loss was used to monitor model overfitting. **B**. Two metrics were used to assess model performance: the F1 score and Confusion Fraction **C**. Macro-averaged class F1 scores for the task of classifying ligand source (host TNF, viral, bacterial, and unstimulated) across polarization states demonstrates loss of stimulus response specificity with polarization **D**. Class F1 scores across polarization states normalized to M0 performance from the same model. **E**. Confusion fractions across polarization states for different ligand sources reveal polarization-dependent patterns in stimulus response specificity. Error bars in C and D and values in E correspond to 95% confidence intervals.

We first applied the LSTM-based classifier to the task of discriminating the ligand sources. To this end we combined NFκB trajectories from Poly(I:C) and R848 under the “viral” label and LPS, Pam3CSK, Flagellin, FSL1 and CpG under the “bacterial” label. We also considered TNF as “host” and had “unstimulated” cell trajectories as well. We found that macrophages showed higher macro-averaged F1 scores in unpolarized naïve conditions than any of the five polarization conditions for the task of classifying different ligand sources, suggesting naïve macrophages have greater stimulus-response specificity than their polarized counterparts (Figure 3C). Naïve macrophages showed the greatest macro-averaged F1 score for the task of classifying each ligand individually as well, and this remained true even when separate models were trained for each polarization state (Figure S3B-D), confirming the loss of stimulus-specificity with polarization. Further, we found that viral ligand identifiability was most diminished by polarization, particularly in IFNβ, IL13, and IL4 polarization states, while host TNF identifiability was the least affected (Figure 3D). We then asked what caused the diminished identifiability by inspecting the confusion fractions between ligand sources (Figure 3E). IL4 polarization resulted in the greatest confusion between both TNF and viral ligands or bacterial ligands. These confusion fractions were also elevated with IL13 polarization, but not with the other M2 polarizer, IL10. IFNβ polarization increased TNF vs. viral ligand confusion only and IFNγ increased TNF vs. bacterial ligand confusion only. All polarization conditions caused convergence of NFκB dynamics in response to viral and bacterial ligand sources. Furthermore, the classifier of individual ligands confirmed these losses of host and pathogen distinguishability with polarization, and revealed convergence among bacterial ligands as well, such as with FSL1 and LPS with interferon polarization (Figure S3E). Finally, confusion with unstimulated conditions was increased for TNF in the IFNβ-polarized state, for viral ligands in the IL13 and IL4 states, and for bacterial ligands in all three states, due to diminished responses in these conditions. Overall, the machine learning analysis revealed losses in the stimulus-specificity of NFκB signaling with all polarizers, but each polarization condition affected different ligand responses differentially.

### Polarizing cytokines have distinct effects on the discrimination of stimuli

We further explored polarization’s effect on the distinguishability of NFκB responses to specific ligands, first examining the increased confusion with IL4 polarization. We now performed classification tasks to identify TNF vs. each of the bacterial ligands (Pam3CSK, Flagellin, CpG, FSL1, and LPS), or TNF vs. each of the viral ligands (R848 and Pol(I:C)). Increased confusion was evident with flagellin, CpG and LPS (Figure 4A), as well as R848 and Poly(I:C) (Figure S4A), thus IL4’s effects were remarkably broad. Deployment of the six signaling codons with TNF and LPS stimulation revealed that LPS responses looked more TNF-like with IL4 polarization due to increased oscillatory content and decreased total activity, reflecting a loss in immune threat (Figure 4B). Similar changes were also apparent with R848 responses (Figure S4C). Exploring the effects of M1 polarizers IFNβ and IFNγ, we found that decreased oscillatory content and increased total activity of TNF responses contributed to the increased confusion with poly(I:C) and flagellin respectively and a gain in immune threat level (Figure S4D-I). Our analysis indicates that the loss of host TNF distinguishability with IL4 polarization is driven by pathogen responses becoming less “pathogen-like”, while for M1 type polarization states, host TNF responses become more “pathogen-like”.

**Figure 4:**
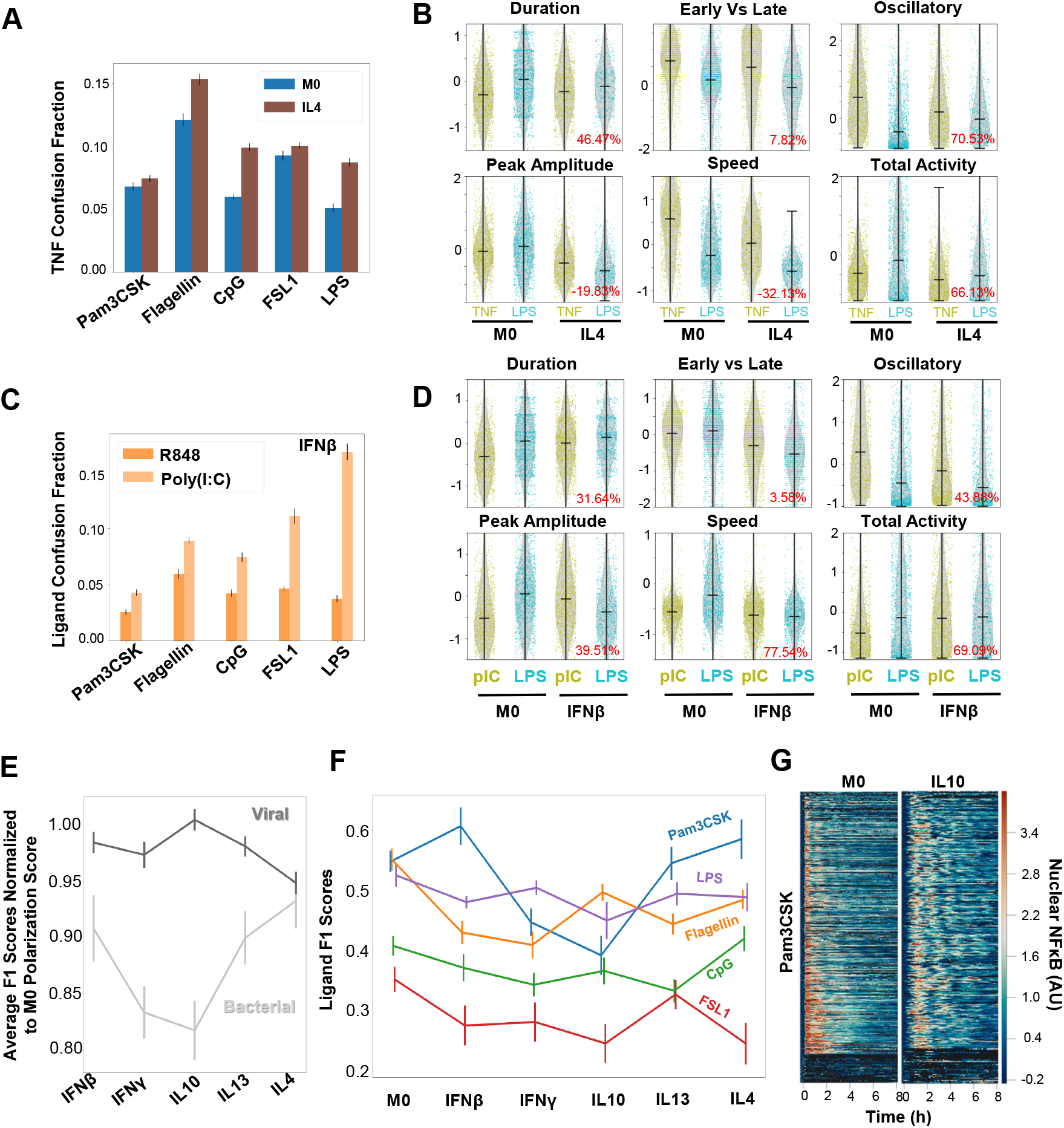
Macrophage polarization affects ligand distinguishability uniquely. **A**. Confusion fraction between the host ligand (TNF) and the bacterial ligands (Pam3CSK, Flagellin, CpG, FSL1, LPS) in the IL4 and M0 polarization states (for the task of individually identifying the host TNF and bacterial ligands) shows larger increase with flagellin, CpG, and LPS stimulation. **B**. Signaling codon distributions from the single-cell responses to TNF and LPS with M0 and IL4 polarization reveal decreased immune threat level of LPS responses (increased oscillations and decreased total activity) drive convergence of stimulus responses with polarization; percent reduction in Jensen-Shannon Distance between ligand responses with polarization in red. **C**. Confusion fraction between the viral ligands (R848, Poly(I:C)) and the bacterial ligands (Pam3CSK, Flagellin, CpG, FSL1, LPS) in the IFNβ polarization state for the task of individually identifying the viral and bacterial ligands; illustrates greatest confusion between Poly(I:C) and LPS. **D**. Signaling codon distributions from the single-cell responses to Poly(I:C) and LPS with M0 and IFNβ polarization reveal an increased immune threat level of both Poly(I:C) (decreased oscillations and increased total activity) and LPS (reduced speed) drives convergence of stimulus responses with polarization. **E**. Macro-averaged class F1 scores normalized to M0 performance from the same model shows greater performance loss for the task of individually classifying the bacterial ligands with polarization compared to that of viral ligands. **F**. Class F1 scores for the task of classifying bacterial ligand responses reveals Pam3CSK as a significant source of confusion with IFNγ and IL10 polarization. **G**. NFκB trajectory datasets with Pam3CSK stimulation in M0 and IL10 polarization demonstrate increased oscillations with polarization. Error bars in A, C, E, and F correspond to 95% confidence intervals.

Next, we investigated further why distinct pathogen response signals converged (Figure 3E), by training a model to individually classify each of the viral vs. bacterial ligands. In the IFNβ polarization state, where viral and bacterial source confusion was the largest, Poly(I:C) showed greatest confusion with LPS, with FSL1 a close second (Figure 4C). We found that all Poly(I:C) and LPS signaling codon distributions became more similar with IFNβ polarization, with the convergence driven most by a diminished oscillatory content and increased total activity in Poly(I:C), and reduced speed in LPS (Figure 4D). These changes correspond to an increase in immune threat characterization for both ligands. We carried out similar analyses in IFNγ, IL10, IL13, and IL4 polarization states (Figure S5); together, these findings suggested that convergence of responses to diverse PAMPs in IFNβ or IFNγ polarization states is due to the generation of a more monolithic or stereotyped “pathogen-like” response signifying a greater threat level, whereas in IL4, IL10, and IL13 they become less “pathogen-like”.

We then examined the ability of macrophages to distinguish particular PAMPs within a pathogen class. We performed two classification tasks: the first discriminated the two viral ligands from each other and the second discriminated the five bacterial ligands from each other. Average F1 scores, normalized to the naïve condition scores, revealed polarization had little effect on viral PAMPs distinguishability, but a big effect on bacterial PAMPs, particularly with IFNγ and IL10 polarization (Figure 4E). Interestingly, each bacterial ligand differentially contributed to this overall classification performance (Figure 4F). Whereas LPS and CpG identifiability was largely unaffected by polarization, Pam3CSK identifiability was diminished most by IFNγ and IL10 polarization, flagellin by both type I and II IFNs, and FSL1 more so by IL10 and IL4.

The ligand-specific effects suggest that different polarization conditions differentially modulate molecular mechanisms that are very proximal to TLRs and MyD88 recruitment. For example, with IL10 polarization there is a significant increase in the oscillatory content of Pam3CSK responses when compared to the naïve condition (Figure 4G), with the mean oscillatory content increasing from −0.18±0.05 to 0.43±0.07. This may be due to a decrease in TLR1/2 surface expression or the recruitment of MyD88. which would diminish downstream IKK activity and hence render the dynamics more oscillatory.

### NFκB stimulus-response dynamics can map macrophage polarization states

Our investigation thus far has focused on how stimulus discrimination based on differences in NFκB response dynamics is affected by macrophage polarization. Our results suggested that polarization states may be distinguishable based on the dynamical NFκB response to a specific stimulus. We used Functional PCA^50^ to dimensionality-reduce the single-cell NFκB trajectories for a specific stimulus, and used the top ten principal components for Uniform Manifold Approximation & Projection (UMAP) to display the 6 polarization states (Figure 5A-B). Visual inspection suggested that TNF stimulation did not reveal much difference between the six polarization states, while Pam3CSK stimulation did. To independently quantify the discrimination of polarization states we trained an LSTM-classifier on polarization conditions.

**Figure 5:**
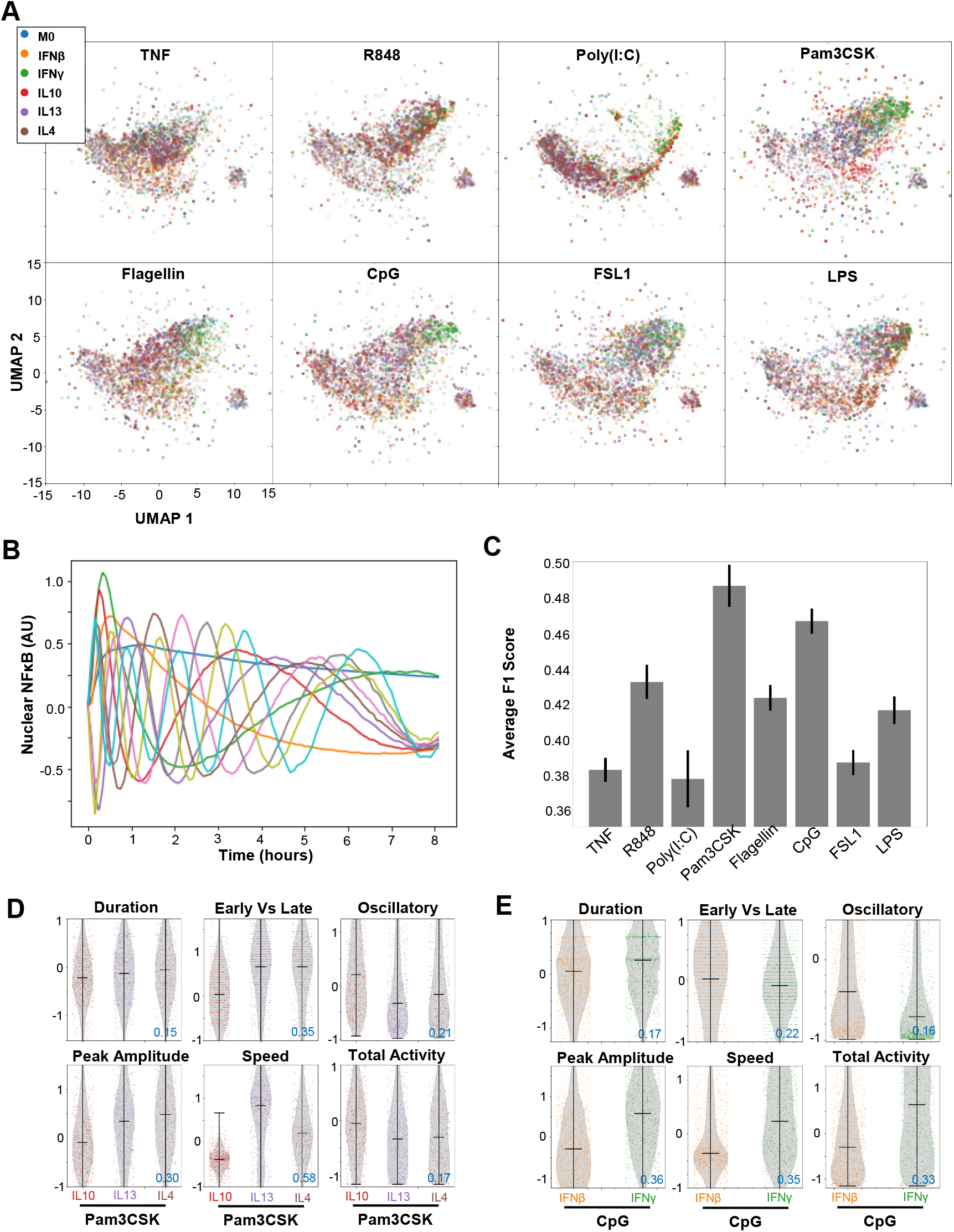
Mapping macrophage polarization states with NFκB signaling response dynamics. **A**. UMAP projection of the first 10 principal components identified by functional PCA (capturing approximately 85.30% of the variance) of the NFκB responses for each stimuli colored by polarization state (down-sampled such that number of samples per condition equivalent) **B**. First 10 principal components used as input for the UMAP projection. **C**. Macro-averaged class F1 scores for the task of classifying each polarization condition provides a quantification of polarizer distinguishability across the stimuli. **D**. Signaling codon distributions from the single-cell responses to Pam3CSK with IL10, IL13, and IL4 polarization reveal separation of IL10 signaling codon values from other M2 polarizers (mean JSD between IL10 and IL13/IL4 in blue). **E**. Signaling codon distributions from the single-cell responses to CpG with IFNβ and IFNγ polarization reveal separation of signaling codon values for the M1 polarizers (JSD between IFNβ and IFNγ in blue).

We found that the classifier had the greatest macro-averaged F1 score with Pam3CSK stimulation, followed by CpG stimulation, while TNF, Poly(I:C), and FSL1 had the least (Figure 5B). Inspecting the UMAP projections, it appears the separation of IL10-polarized cells from other M2-polarized cells contributes to the increased performance with Pam3CSK, while some separation of IFNβ- and IFNγ-polarized cells is relevant in the case of CpG stimulation.

Examining the signaling codons of the NFκB dynamics in response to these stimuli illustrates how polarizers alter dynamics, particularly amongst M1 and M2 type polarizers. IL10-polarized responses to Pam3CSK differ from those polarized with IL13 and IL4 due to their less early activity, peak amplitude, and speed, as well as more oscillations (Figure 5D). IFNβ-polarized responses to CpG differ from those polarized with IFNγ due to their peak amplitude, speed, and total activity, as well as more early activity (Figure 5E). Overall, these findings illustrate that macrophage polarization states cannot be realistically described along a single dimension, but form a multi-dimensional landscape.

This analysis highlights that the ability to discriminate polarization states based on NFκB response dynamics depends on the stimulus used to elicit NFκB activation. Presumably better discrimination of polarization states could be achieved if NFκB dynamics in response to multiple stimuli were available for each cell. However, each cell can be interrogated experimentally only by a single stimulus. To integrate signaling dynamic information from all stimuli, future studies may involve a mathematical model of the NFκB signaling network with parameter distributions inferred from the experimental data to undertake *in silico* simulations of single macrophages responding to different stimuli.

## DISCUSSION

Macrophages are subjected to diverse tissue microenvironments characterized by a variety of cytokines that modulate their functions. One functional hallmark of macrophages is their ability to mount stimulus-specific immune responses to diverse immune threats, as observed in studies of gene expression^18,19,51^ or NFκB dynamics^30^. Here, we explored the effect of macrophage polarization on stimulus-specific NFκB response dynamics by generating an unprecedented dataset of single-cell NFκB response trajectories associated with a wide array of polarizing cytokines and stimulating ligands, and developing analytical frameworks for interpreting these datasets. Our analysis revealed polarization-specific effects on NFκB dynamics and response specificity that could be traced to the response to specific stimuli and specific dynamic features. Our results suggest how macrophage polarization states do not lie on a continuum from M1 to M2 states, but rather a landscape of distinct functional specialization states.

Prior work has aimed to identify transcriptional and epigenetic signatures of macrophage states^52–58^. More recently, single-cell sequencing approaches have been leveraged for an unbiased data-driven characterization of macrophage states in various physiologic and pathologic contexts^59–64^. These studies rely on profiling the abundance of molecules using a snap-shot measurement to characterize macrophage states. In contrast, our study probes the functionality of the macrophage. The function we are able to probe here at the single cell level is the stimulus-response signaling dynamics of NFκB. This function does not linearly result from the chromatin landscape or mRNA abundances but involves non-linear assemblies of protein complexes, membrane organization, and transport processes. As demonstrated here, NFκB signaling dynamics can be leveraged to capture this additional information enabling an alternative mapping of polarization states based not on snap-shot transcriptomic or epigenomic data but on a functional biological response.

Developing appropriate computational tools to measure the distinguishability between different single-cell NFκB trajectories was essential to address several analysis challenges that arise from working with large time-series datasets. To determine distinction between trajectories, a reliable notion of distance must be established; however, for time-series data standard definitions of distance (i.e. Euclidean distance) often fail to capture differences appropriately. Specifically, treating a n-timepoint trajectory as a n-dimensional vector disregards the relationship between timepoints, and two single-cell trajectories with similar dynamical patterns but slightly displaced in time could be computed to be highly distinct^65^. To describe differences between experimental conditions, summary statistics are easy to compute and interpret, however they are insufficient. For example, taking the timepoint by timepoint mean of the single-cell trajectories can obscure asynchronous oscillatory dynamics observed at the single-cell level^66^. Further, average behavior descriptions mask how the distributions of responses actually overlap. Employing measures of spread or shape that are used to characterize distributions, are also not fully informative if taken at a timepoint level, because they also do not recognize the inter-timepoint correlations and so risk overestimating the dispersion.

We addressed these challenges using two innovations. The first is the utilization of ‘signaling codons’, which are features of dynamical trajectories that are informative about the stimulus, as defined by mutual information^30^. Thus, six signaling codons sufficiently describe the stimulus-specific dynamical NFκB trajectories and their values are more robust to temporal shifts previously discussed. In essence, they constitute a lower dimensional representation of the data, thereby expanding the range of analysis tools that can be used, while preserving biological interpretability. Our second approach to address the challenges of time-series data analysis was to utilize a novel machine learning approach that allowed for trajectory distinguishability to be explored in a feature-free manner. Indeed, the LSTM classifier performs with higher precision than a standard ensemble of decision tree classifier trained with time-series data. The LSTM architecture enabled direct analysis of the time-series data which allows for the recognition of informative variation not limited by predefined features. This approach also permits an interrogation of distinguishability that includes the single-cell resolution.

We first used the NFκB signaling codons in a LDA analysis that was based on the concept that NFκB responses encode immune threat level of the encountered stimuli with the projection maximizing the distinction between the responses to PAMPs and host TNF. This data-driven approach ordered each paired PAMP/polarization condition along a continuum and revealed that the immune threat character of responses was maximized in M1 polarization states mediated by interferons and minimized in M2 polarization states mediated by IL4, IL10, and IL13. The approach also revealed which signaling codons were associated with an increased immune threat response: larger peak amplitude and total activity, and fewer oscillations and less speed. Furthermore, we utilized this descriptive framework to interpret the signaling codon changes that drove convergence with polarization. Broadly, elevated immune threat responses contributed to convergence in M1 polarizations states, whereas the opposite was observed for M2 polarization states. Hence, the immune threat level axis can be associated with the pro-vs anti-inflammatory M1/M2 dichotomy and a continuum of polarization states between these poles elicited by various microenvironmental cytokines^3,67,68^.

LSTM-based ML analysis provided additional insights. It identified increased confusion between IL4-treated macrophage responses to TNF and pathogen ligands, mediated by responses to pathogens, such as R848 and LPS, having more oscillatory but less total activity with polarization, reducing immune threat with this M2 type polarization. In contrast, the convergence of TNF and pathogen responses with M1 polarization states was driven by opposite changes of the same signaling codons in TNF, elevating immune threat level. We additionally detected increased confusion between the viral and bacterial ligand sources across all polarization states, suggesting a convergence of pathogen responses with polarization. We discovered, however, that this convergence was mediated by pathogen responses attaining greater immune threat characteristics with M1 polarization conditions and losing them with M2 polarization.

While these results could be related to the conception of a linear continuum in M1 vs M2 polarization states, our analysis also revealed its limitations as we found for example differences within these M1 and M2 polarization states that are specific to particular polarizing cytokines.

Firstly, the immune threat characterization of ligands varied between IFNβ- and IFNγ-treated macrophages, as well as between IL10, IL13, and IL4 treated macrophages. For example, for the viral ligand R848, the immune threat level of its responses was elevated with IFNγ polarization, but slightly diminished with IFNβ polarization. For the bacterial ligand Pam3CSK, the immune threat level of its responses was significantly minimized with IL13 polarization, but much less so with IL10 and IL4 polarization. Furthermore, there were polarization-specific effects to losses in NFκB response distinguishability. For example, within the M2 polarizers, IL4 led to the greatest confusion between host TNF and pathogen responses as well as between viral and bacterial source responses, however IL10 led to the greatest confusion between different bacterial ligands. These areas of confusion were mediated by similar signaling codon changes, notably increased oscillations of pathogen associated responses; however, these changes manifested differently in the ligand responses with each polarization state, hence resulting in distinguishability differences. Finally, projecting the NFκB response dynamics to the different stimuli using functional PCA revealed a mapping of the polarization states that provided separation between M1 polarizers from one another, as well as M2 polarizers from one another in certain stimulation conditions, like Pam3CSK and CpG respectively. These effects suggest a conception of macrophage polarization that occupies a higher number of dimensions as there are several axes along which these states can differ from one another, going beyond the M1/M2 continuum.

These findings support the notion of finer functional niches within macrophage activation. Macrophages stimulated by IL4 versus those stimulated with IL10 for example have been previously identified as functionally distinct, with the former serving a more wound-healing role and the latter serving a more regulatory role and hence differences in stimulus-response specificity may support these functions^69^. M2a macrophages have been associated with increased susceptibility to viral, bacterial, and fungal infections which may align with poor host versus pathogen recognition. M2c macrophages are associated with late stages of adaptive immune response and dampening the response, and hence the ability to differentiate bacterial pathogens may be nonessential. Such functional characterization of macrophage subtypes has been previously studied in atherosclerotic and dermatological lesions which identified macrophage subtypes beyond the M1/M2 dichotomy *in vivo*^70,71^. Our study supports the notion that macrophage activation is better described as a multi-dimensional topology with different functional zones rather than a linear continuum between two functional poles. In that sense the process of macrophage polarization can then perhaps be best analogized with the concept of a “Waddington landscape”. Naïve macrophages at the outset have tremendous specialization potential and upon microenvironmental exposures the functional capacities of the macrophage narrow. This specialization of macrophage function aligns with the losses in response specificity that we observed with polarization. Unlike naïve macrophages that must maintain a high degree of functional pleiotropy, polarized macrophages have less need to distinguish stimuli in their prescribed effector roles, and hence a canalization of stimuli responses is appropriate. Future studies may further describe this functional landscape of macrophage polarization states and the transition between them, by combining our signaling data with other single-cell measurements or using mathematical models of the signaling network that account for the observed signaling dynamics^72^.

## ACKNOWLEDGEMENTS

Authors acknowledge Roberto Spreafico for contributing to the establishment of the experimental cell system. A.S. was supported by 2T32GM008185 and 2T32GM008042. Funding was provided by R01AI127864, R01AI32835, R01AI127867 to AH.

## AUTHOR CONTRIBUTIONS

Conceptualization, A.H.; Methodology, S.S. (data generation), A.S. (data analysis, modeling, interpretation), A.A. (establishing workflows); Investigation, S.S., A.S., A.A.; Writing – Original Draft, A.S., A.H., Writing – Review & Editing, XXXXX. Supervision, Project Administration, Funding Acquisition, A.H.

## INCLUSION AND DIVERSITY STATEMENT

One or more of the authors of this paper self-identifies as an underrepresented ethnic minority and received support from a program designed to increase minority representation in science. While citing references scientifically relevant for this work, we actively worked to promote gender balance in our reference list.

## RESOURCE AVAILABILITY

### Lead contact and materials availability

Further information and requests for resources and reagents should be directed to and will be fulfilled by the lead contact, Alexander Hoffmann.

### Data and code availability

Single-cell RNA-seq data have been deposited at SRA under BioProject accession number PRJNA819468 and are publicly available as of the date of publication. Accession numbers are listed in the key resources table. Trajectory data generated from microscopy experiments have been deposited at Mendeley Data (https://doi.org/10.17632/gkxzb5hcmk.1) and are publicly available as of the date of publication. Software for image analysis is available on GitHub (https://github.com/brookstaylorjr/MACKtrack) and code to calculate signaling codons is available on GitHub (https://github.com/signalingsystemslab) as of the date of publication.

## EXPERIMENTAL MODEL AND SUBJECT DETAILS

### Macrophage Cell Culture and Stimulation

Immortalized myeloid precursor (iMP) cells were prepared from RelA-mVenus mouse strain^30^ by HoxB4-mediated transduction ^43^. iMP-Derived Macrophages (iMPDMs) were prepared by culturing iMPs in L929-conditioned medium using standard Bone-Marrow Derived Macrophage (BMDM) culture method ^30^ iMPDMs were re-plated in imaging dishes on day 6 at 20,000 cells/well in an 8-well ibidi SlideTek chamber, for imaging at an appropriate density on day 10 or day 11. iMPDMs were treated with polarization reagents (IL4 (10 ng/mL), IL13 (50 ng/mL), IL10 (20 ng/ML) IFNγ (10 ng/mL) or IFNβ (100 U/ML)) 24 hrs before stimulation. Stimulation was done with the toll-like receptor (TLR) 4 agonist, lipopolysaccharide (LPS) (Sigma Aldrich), TLR3 agonist, polyinosine-polycytidylic acid (Poly(I:C) (Invivogen), TLR9 agonist, CpG B ODN (invivogen); TLR2 agonists, Pam3CSK4 (invivogen) and FSL1, TLR8 agonist, R848 (invivogen) or cytokine TNF (R&D Systems) without media replacement.

## METHOD DETAILS

### RNA Isolation and Sequencing

Bone-Marrow Derived Macrophages (BMDMs) were cultured with standard methods, L929-conditioned medium^30^. Raw 264.7 cells were cultured in DMEM 10 % FBS media. After stimulation, cells were harvested at desired time points. For PolyA+ RNA, cells were harvested in TRIzol reagent (Life Technologies, Carlsbad, CA). Then, DNA-free RNA was extracted from cell using DIRECTzol kit (Zymo Research, Irvine, CA) according to manufacturer’s instructions. After RNA extraction, libraries for polyA+ RNA were prepared using KAPA Stranded RNA-Seq Kit for Illumina Platforms (KAPA Biosystems, Wilmington, MA) according to the manufacturer’s instructions. Resulting cDNA libraries were single-end sequenced with a length of 50bp on an Illumina HiSeq 2000 (Illumina, San Diego, CA).

### Sequencing Mapping and Analysis of RNA-Seq

After adapter trimming with cutadapt^73^, sequences were preprocessed with PRINSEQ^74^ using the “dust” method to filter low complexity sequences with the maximum allowed score set to 7 and sequences with more than 10% ambiguous bases were removed. Single-end reads were mapped to reference mouse genome (mm10) using STAR^75^ with the following options: -- outFilterMultimapNmax 20 --alignSJoverhangMin 8 --alignSJDBoverhangMin 1 -- outFilterMismatchNmax 999 --outFilterMismatchNoverLmax 0.04 --alignIntronMin 20 -- alignIntronMax 1000000 --alignMatesGapMax 1000000 --seedSearchStartLmax 30. Only primary mapped reads with alignment score (MAPQ)>30 were then selected by Samtools^76^. Ribosomal RNA was filtered out using the intersect function in bedtools with a minimal overlap fraction of 0.1 and finally reads mapped to the Y chromosome or mitochondria were removed for downstream analysis. Transcript abundance was quantified based on GENECODE M4 annotation using featureCounts^77^ using option ‘-t exon -g gene_id. For analysis, genes with no count across all experiments were filtered out. An average pseudocount of 2 was added to the raw counts, where the exact value added to each library was proportional to the library size.

The counts were then normalized for differences in library size by calculating the counts per million (CPM) and then the base 2 log of those values were used to calculate fold change. Genes induced by LPS were determined to be those that had a log2 Fold Change greater than or equal to 1 after 3 hours post LPS stimulation in two replicate experiments of BMDM’s.

### Live-cell imaging

2 hours prior to imaging, iMPDMs were stained with nuclear staining dye, Hoechst 33342 (5 ng/mL). ibidi chamber was placed to imaging station. Cells were imaged at 5-minute intervals on a Zeiss AxioObserver platform with live-cell incubation, using epifluorescent excitation from a Sutter Lambda XL light source. The first three images collected (pre-stimulation) were used to determine the baseline activity of NFκB for each cell. After 15 mins of the start of imaging, conditioned culture media containing stimulus was injected into the respective well of ibidi chamber *in situ*. Images were recorded on a Hamamatsu Orca Flash 2.0 CCD camera for 12.5 hrs

## QUANTIFICATION AND STATISTICAL ANALYSIS

### Image analysis and processing

Microscopy time-lapse images were exported for single-cell tracking and measurement in MATLAB R2018a, used in earlier work^30^. Briefly, cells were identified using DIC images, then segmented, guided by nuclear staining from the Hoechst image. Segmented cells were linked into trajectories across successive images, then nuclear and cytoplasmic boundaries were defined and used for measurement in fluorescent channel for mVenus-NFκB. Nuclear NFκB levels were quantified on a per-cell basis, normalized to image background levels, then were baseline-subtracted. The first three images collected (pre-stimulation) were used to determine the baseline activity of NFκB for each cell. The mean fluorescence value from these three frames was subtracted from the complete trajectory to normalize each cell. For downstream analysis and visualization, the third timepoint corresponds to time = 0 and 97 timepoints after that were included (∼ 8 hour trajectories). Mitotic cells, as well as cells that drifted out of the field of view, were excluded from analysis. The code (MACKtrack) used for this analysis are publicly available at GitHub (https://github.com/brookstaylorjr/MACKtrack).

### Signaling Codon Calculations

To quantify the 6 signaling codons, 11 metrics were applied to the NFκB trajectories (Table S1). For signaling codons formed by more than one trajectory feature, the trajectory features were z-scored and the mean of the z-scores was taken to get the signaling codon value. During quality control analysis to determine biological replicates, z-scoring was performed over cells in the experimental condition of interest. Additionally, for the quality control analysis, the trajectory features from only “responding” cells were considered. A cell was deemed a responder if its trajectory exceeded three times the standard deviation of the baseline for at least 5 consecutive time points. Experiments were finally deemed biological replicates if the Jensen-Shannon distance (JSD) between each of their signaling codon distributions were below a pre-specified threshold, 0.3. For all subsequent analysis and visualizations presented in this paper, z-scoring was performed over all cells in all experimental conditions listed in Figure 1B to calculate signaling codon values. The code to calculate these signaling codons values are provided on the GitHub site.

To calculate the Jensen-Shannon distances (JSD) between signaling codon distributions, the Freedman-Diaconis rule^78^ was first used to select a bin width for each signaling codon. Using this bin width and the extremum signaling codon values, a histogram that approximates the probability density function for each experiment can be constructed and used to calculate the JSD (the square root of the Jensen Shannon Divergence using the base 2 logarithm). For LDA and PCA calculations using the signaling codons, the scikit-learn Python package^79^ was used, cells with missing values were excluded, and the standard scaler was applied. The confidence intervals reported were two-sided and used a normal distribution with associated z-scores.

### LSTM-based Machine Learning Classifier

The LSTM-based Machine Learning Classifier was implemented in TensorFlow 2^80^ using the Keras API^81^. The classifier utilized the trajectories from time = 0 to 8.083 hours for a total of 98 timepoints. Trajectories with missing (nan) values were excluded from this analysis. For each classification task described, trajectories were sampled from each polarization state such that for each combination of class and polarization state the number of trajectories were equivalent. More specifically, for each combination of class and polarization state the trajectories were either undersampled or resampled to reach the mean number of trajectories across the class and polarization state combinations. The sampled data was then split 60% for training, 20% for validation, and 20% for testing. For each classification task, the data was shuffled and resplit 15 times to estimate uncertainty in output performance metrics. The confidence intervals reported were two-sided and used a T-distribution with degrees of freedom one less than the sample size (n-1). A standard scaling, fit from the training data, was finally applied across each time point.

The architecture of the machine learning classifier consisted of a LSTM layer with the dimensionality of the output set to the number of timepoints, 98, followed by a fully connected layer with the dimensionality of the output set to the number of classes. A softmax activation function was finally applied to the output of the fully connected layer. The weights of the classifier were optimized by minimizing the categorical cross-entropy loss objective function with the Adam algorithm using the following default parameters: learning rate=0.001, beta 1=0.9, beta 2=0.99, epsilon = 1e-08, batch size=32. With increasing number of training epochs, the value of the loss function over the training data will continue to decrease whereas eventually the value of the loss function over the validation data (data unseen during optimization) will begin to increase. This signals overfitting, as the trained model loses generalizability of its performance on new data. We employed a simple early stopping technique to address this. For each classification task, the validation loss was monitored during training and the epoch number corresponding approximately to the start of the rise in validation loss was determined. Training was then terminated just prior to this epoch (typically around 60-80 epochs of training in total).

The testing data held out during training was finally used to evaluate the performance of the trained model. The output of the classifier is the probability that a trajectory belongs to each class. To assign the trajectory to a class, the class with the highest prediction probability for each trajectory gave the assignment. These output prediction probabilities and class assignments from the testing data were then used to calculate the performance metrics as described.

### Functional Principal Component Analysis

Functional principal component analysis of the NFκB response trajectories across all stimulation and polarization conditions was performed using scikit-fda^82^. An equal number of samples from each experimental condition was used. This analysis operated directly on the centered raw data (*discretized FPCA*) without first converting the data using a basis representation. The first ten principal components were then utilized to create a UMAP projection of the data using the Uniform Manifold Approximation & Projection package^83^ with default parameters.

---

**Figure S1:**
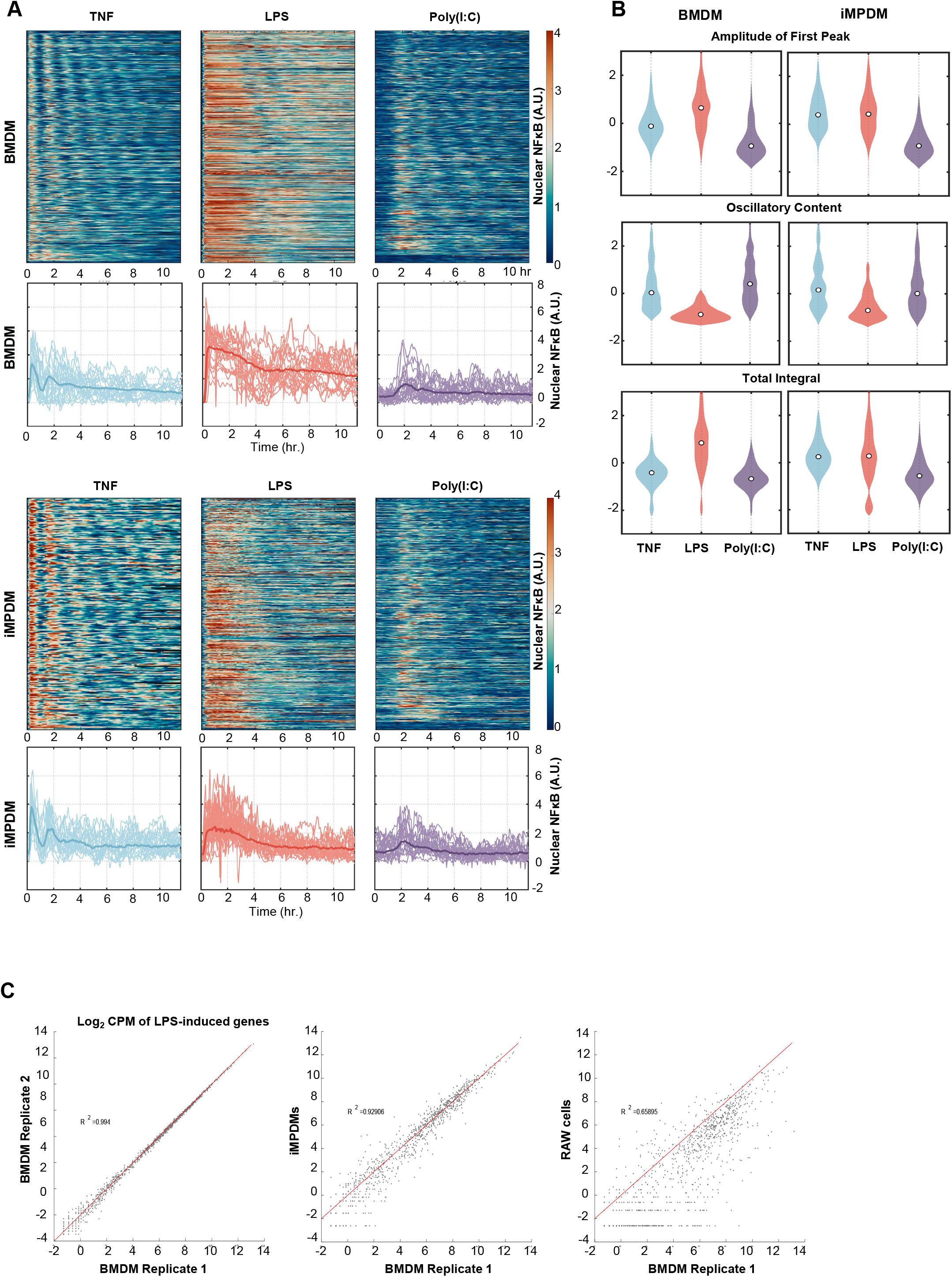
Stimulus-responsive NFκB signaling dynamics and gene expression in iMPDMs (immortalized myeloid progenitor derived macrophages) and BMDMs (bone-marrow derived macrophages) are similar. **A**. Heatmaps of single-cell NFκB trajectories in response to stimulation with TNF, LPS, and Poly(I:C) produced in BMDMs (top), and iMPDMs (bottom) **B**. Distribution of normalized NFκB trajectory features in BMDM and iMPDM single cell responses to TNF, LPS, and Poly(I:C) stimulation **C**. Log2 CPM of gene expression following 3 hours of LPS stimulation in BMDM, iMPDM, and RAW cells. LPS-induced genes are defined as having a Log2 Fold Change equal to or greater than 1 compared to unstimulated basal expression in two replicates of BMDMs.

**Figure S2:**
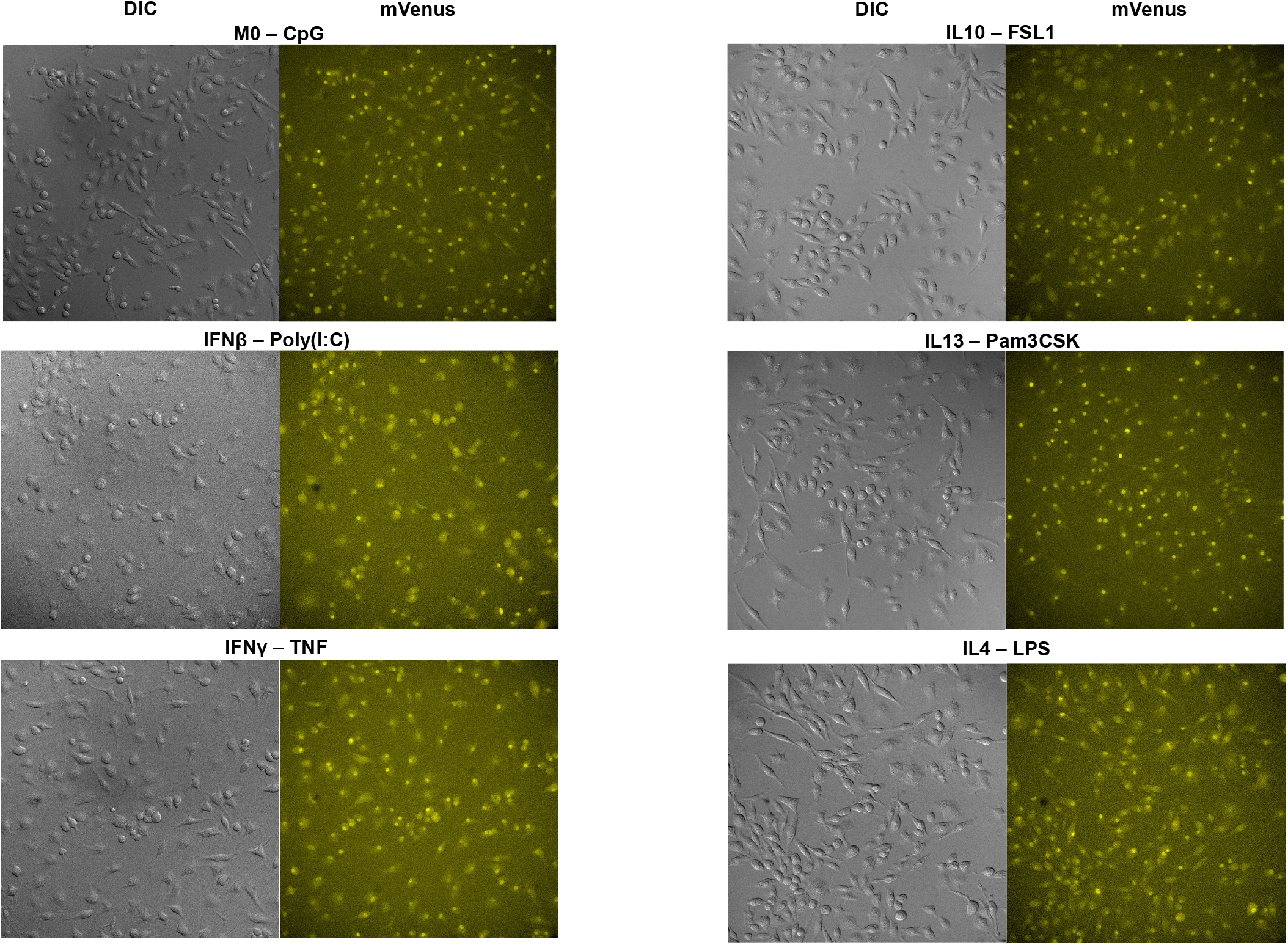
Representative brightfield and fluorescence microscopy images demonstrate that iMPDMs (immortalized myeloid progenitor derived macrophages) appear healthy under different polarizing conditions and mVenus-RelA localizes to the nucleus across the various polarization and stimulation conditions.

**Figure S3:**
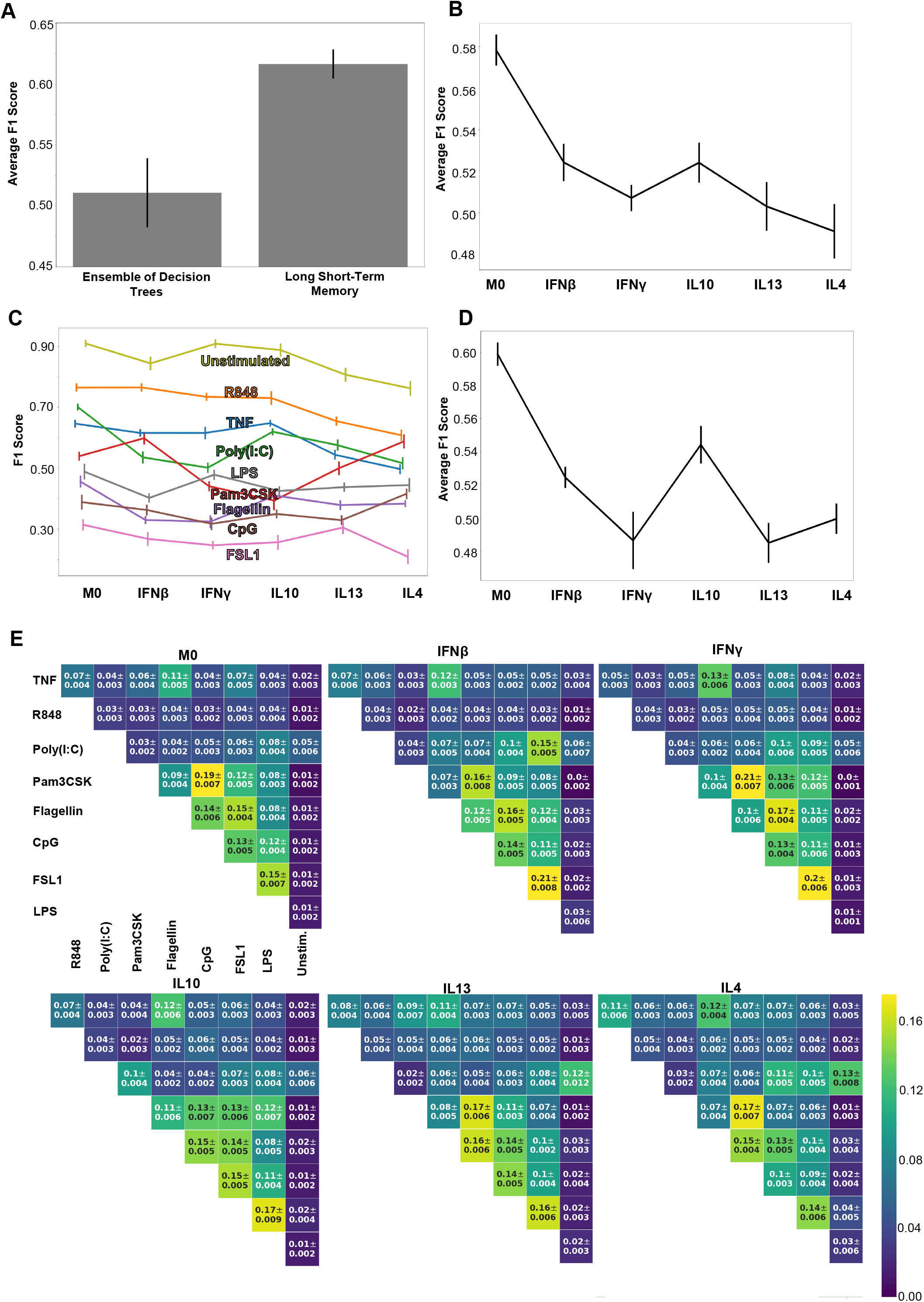
LSTM-based ML classifier performance. **A**. Comparison of the macro-averaged F1 scores for the task of identifying each ligand 0.00 from the time series data in the naïve condition using the ensemble of decision trees versus the LSTM-based classifier. **B**. Macro-averaged class F1 scores for the task of classifying each ligand individually across all polarization states reveal overall loss of specificity with polarization **C**. Class F1 scores across polarization states for the same task as in B. **D**. Macro-averaged class F1 scores for the task of classifying each ligand individually with a model trained separately for each polarization state again reveals overall loss of specificity with polarization. **E**. Confusion fractions across polarization states for different ligand stimulations demonstrate polarization specific patterns in stimulus response specificity. Error bars in A, B, C, & D and values in E correspond to 95% confidence intervals.

**Figure S4:**
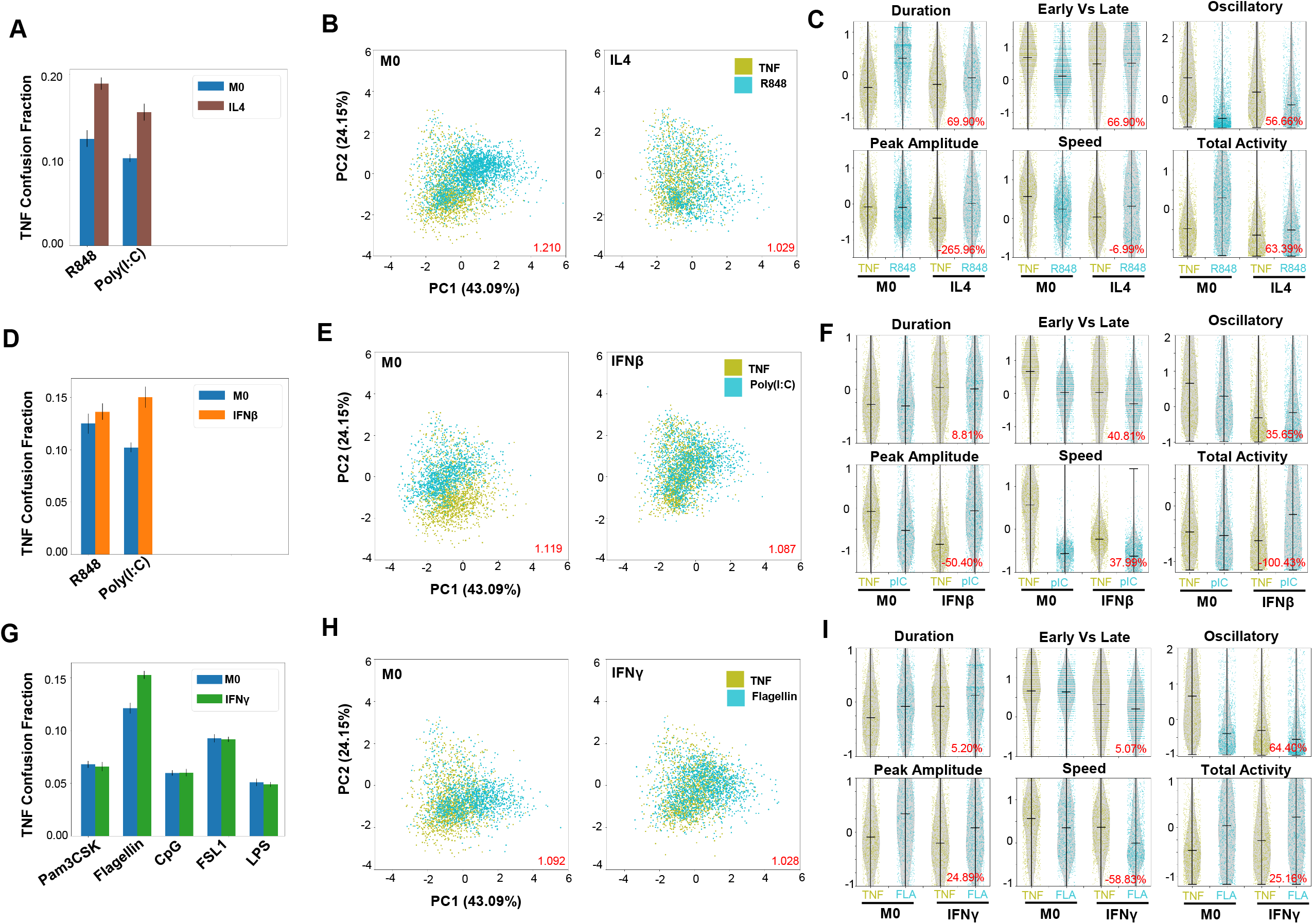
Additional examples of increased host TNF confusion with polarization. **A**. Confusion fraction between the host ligand (TNF) and the viral ligands (R848 and Poly(I:C)) in the IL4 and M0 polarization states for the task of individually identifying the host TNF and viral ligands shows greater increase with R848 stimulation. **B**. PCA projection of the signaling codons from the single-cell responses to TNF and R848 with M0 and IL4 polarization; dispersion measure in red (average pairwise distance between classes divided by average pairwise distance within classes) illustrates convergence of stimulus responses with IL4 polarization **C**. Signaling codon distributions from the single-cell responses to TNF and R848 with M0 and IL4 polarization reveal a decreased immune threat level of R848 responses (increased oscillations and decreased total activity) drive convergence; percent reduction in Jensen-Shannon Distance between ligand responses with polarization in red. **D**. Confusion fraction between the host ligand (TNF) and the viral ligands (R848, Poly(I:C)) in the M0 and IFNβ polarization states shows increased confusion between Poly(I:C) and TNF. **E**. PCA projection of the signaling codons from the single-cell responses to TNF and Poly(I:C) with M0 and IFNβ polarization; dispersion measure illustrates convergence of stimulus responses **F**. Signaling codon distributions from the single-cell responses to TNF and Poly(I:C) (pIC) with M0 and IFNβ polarization reveal an increased immune threat level of TNF responses (decreased oscillations and increased total activity) drive convergence. **G**. Confusion fraction between the host ligand (TNF) and the bacterial ligands (Pam3CSK, Flagellin, CpG, FSL1, LPS) in the M0 and IFNγ polarization states show greatest increase with Flagellin stimulation. **H**. PCA projection of the signaling codons from the single-cell responses to TNF and Flagellin with M0 and IFNγ polarization; dispersion measure illustrates convergence of stimulus responses. **G**. Signaling codon distributions from the single-cell responses to TNF and Flagellin (FLA) with M0 and IFNγ polarization reveal an increased immune threat level of TNF responses (decreased oscillations and increased total activity) drive convergence. Error bars in A and D correspond to 95% confidence intervals.

**Figure S5:**
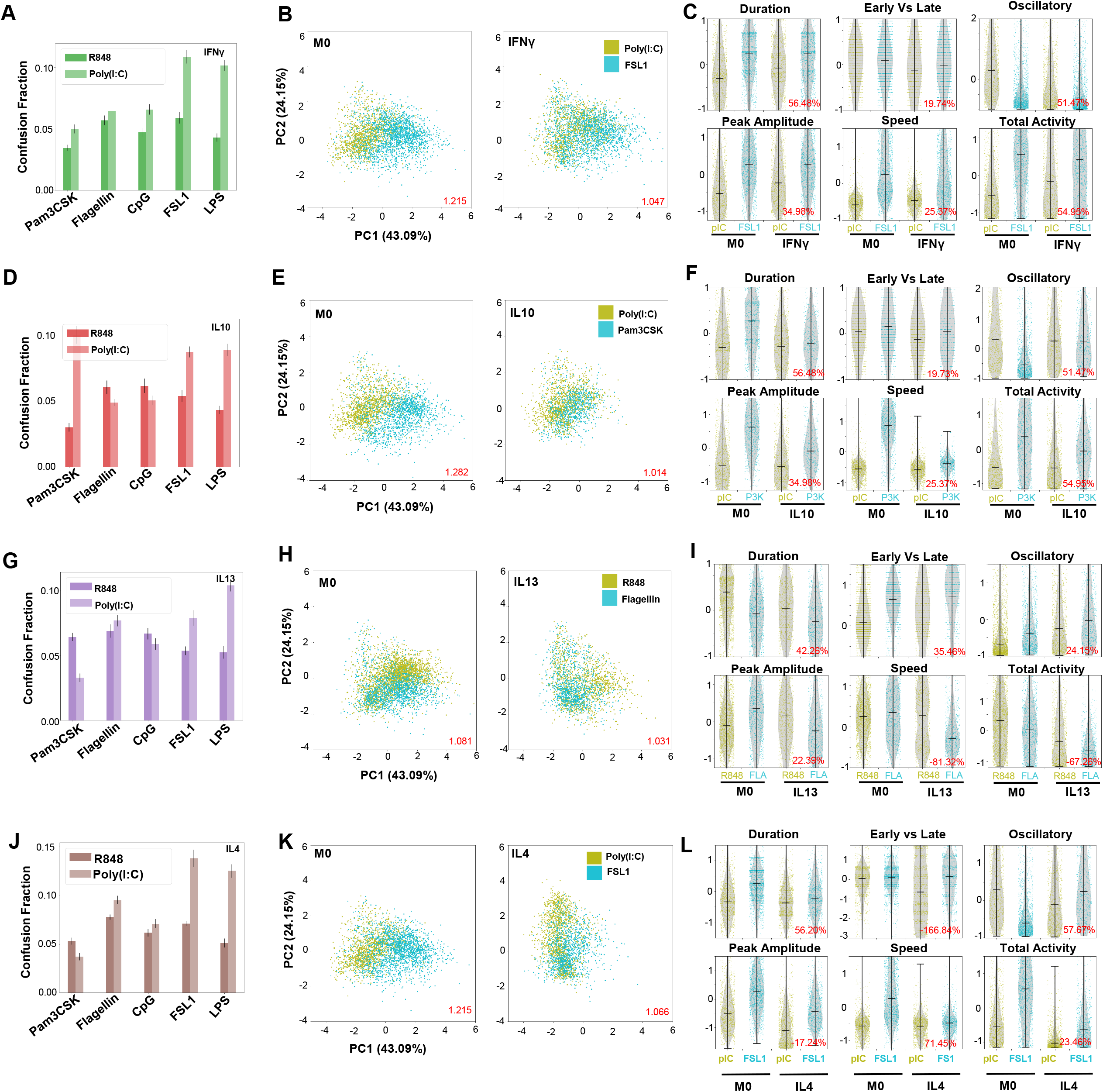
Additional examples of the convergence of viral and bacterial responses with macrophage polarization. **A**. Confusion fraction between the viral ligands (R848, Poly(I:C)) and the bacterial ligands (Pam3CSK, Flagellin, CpG, FSL1, LPS) in the M1 IFNγ polarization state identified elevated confusion between Poly(I:C) and FSL responses **B**. PCA projection of the signaling codons from the single-cell responses to Poly(I:C) and FSL1 with M0 and IFNγ polarization; dispersion measure in red (average pairwise distance between classes divided by average pairwise distance within classes) shows convergence of stimulus responses with polarization **C**. Signaling codon distributions from the single-cell responses to Poly(I:C) (pIC) and FSL1 with M0 and IFNγ polarization reveal increased immune threat of Poly(I:C) (increased duration, peak amplitude, & total activity and decreased oscillations) drives convergence; percent reduction in Jensen-Shannon Distance between ligand responses with polarization in red. **D**. Confusion fraction between the viral and the bacterial ligands in the M2 IL10 polarization state identified elevated confusion between Poly(I:C) and Pam3CSK. **E**. PCA projection of the signaling codons from the single-cell responses to Poly(I:C) and Pam3CSK with M0 and IL10 polarization shows convergence of stimulus responses with polarization. **F**. Signaling codon distributions from the single-cell responses to Poly(I:C) (pIC) and Pam3CSK (P3K) with M0 and IL10 polarization reveal decreased immune threat level of Pam3CSK (decreased duration, peak amplitude, & total activity and increased oscillations) drives convergence. **G**. Confusion fraction between the viral and bacterial ligands in the M2 IL13 polarization state identified Flagellin responses as most confused with R848 responses. **H**. PCA projection of the signaling codons from the single-cell responses to R848 and Flagellin with M0 and IL13 polarization shows convergence of stimulus responses with polarization. **I**. Signaling codon distributions from the single-cell responses to R848 and Flagellin (FLA) with M0 and IL13 polarization reveal decreased immune threat level of R848 (decreased duration and increased oscillations) drive convergence. **J**. Confusion fraction between the viral and bacterial ligands in the M2 IL4 polarization state identified elevated confusion between Poly(I:C) and FSL1. **H**. PCA projection of the signaling codons from the single-cell responses to Poly(I:C) and FSL1 with M0 and IL4 polarization shows convergence of stimulus responses with polarization. **L**. Signaling codon distributions from the single-cell responses to Poly(I:C) and FSL1 with M0 and IL4 polarization reveal decreased immune threat level of FSL1 (decreased duration & total activity and increased oscillations) drives convergence. Error bars in A, D, G and J correspond to 95% confidence intervals.

**Table S1:**
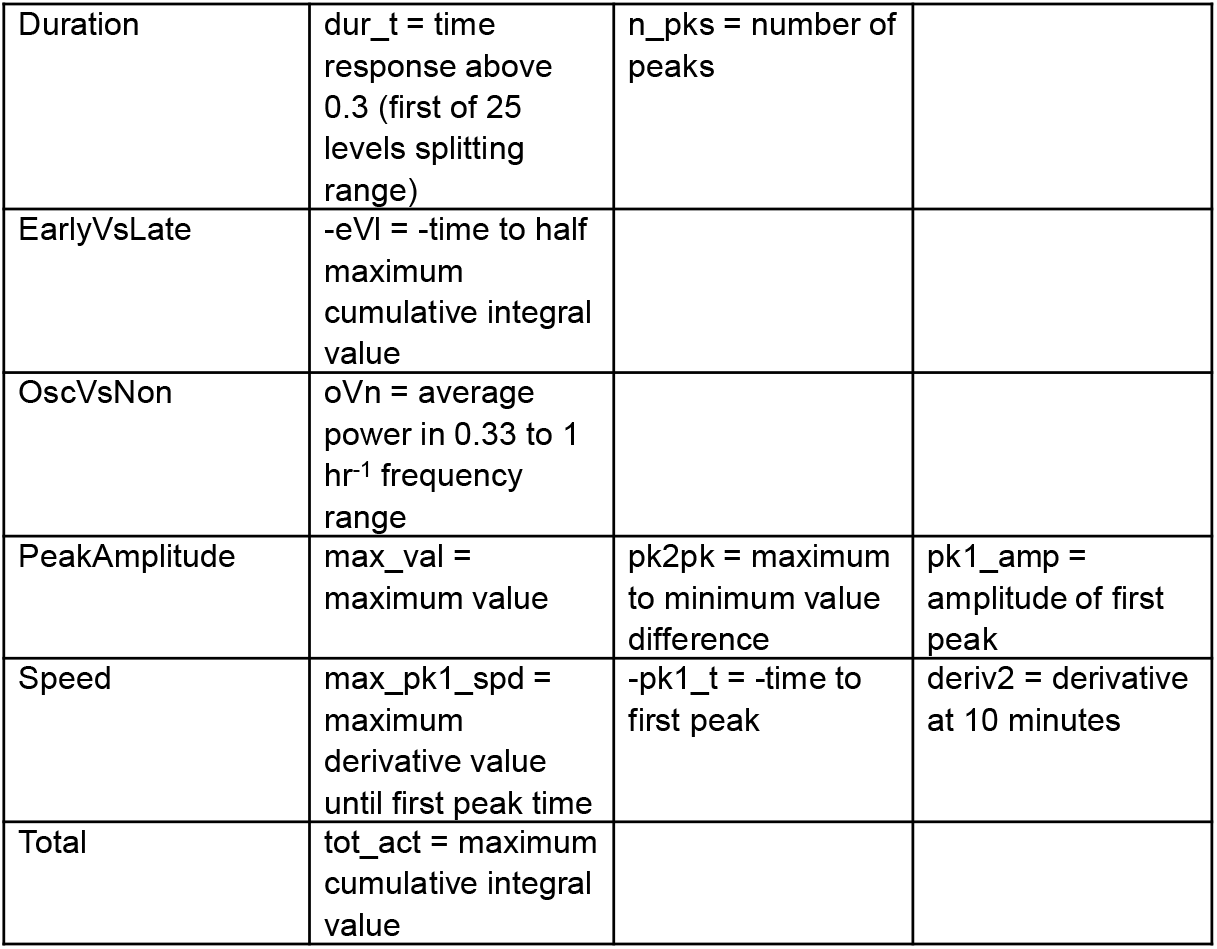
Signaling Codon Definitions.

|  |  |  |  |
| --- | --- | --- | --- |
| Duration | dur_t = time response above 0.3 (first of 25 levels splitting range) | n_pks = number of peaks |  |
| EarlyVsLate | -eVI = -time to half maximum cumulative integral value |  |  |
| OscVsNon | oVn = average power in 0.33 to 1 hr<sup>-1</sup> frequency range |  |  |
| PeakAmplitude | max_val = maximum value | pk2pk = maximum to minimum value difference | pk1_amp = amplitude of first peak |
| Speed | max_pk1_spd = maximum derivative value until first peak time | -pk1_t = -time to first peak | deriv2 = derivative at 10 minutes |
| Total | tot_act = maximum cumulative integral value |  |  |

## REERENCES

1. Murray PJ, Wynn TA. Protective and pathogenic functions of macrophage subsets. Nat Rev Immunol. 2011;11(11):723–737. doi:10.1038/nri3073

2. Murray PJ. Macrophage Polarization. Annu Rev Physiol. 2017;79(1):541–566. doi:10.1146/annurev-physiol-022516-034339

3. Atri C, Guerfali FZ, Laouini D. Role of Human Macrophage Polarization in Inflammation during Infectious Diseases. Int J Mol Sci. 2018;19(6):1801. doi:10.3390/ijms19061801

4. Wynn TA, Chawla A, Pollard JW. Macrophage biology in development, homeostasis and disease. Nature. 2013;496(7446):445–455. doi:10.1038/nature12034

5. Martinez FO, Sica A, Mantovani A, Locati M. Macrophage activation and polarization. Front Biosci J Virtual Libr. 2008;13:453–461. doi:10.2741/2692

6. Sica A, Erreni M, Allavena P, Porta C. Macrophage polarization in pathology. Cell Mol Life Sci. 2015;72(21):4111–4126. doi:10.1007/s00018-015-1995-y

7. Funes SC, Rios M, Escobar-Vera J, Kalergis AM. Implications of macrophage polarization in autoimmunity. Immunology. 2018;154(2):186–195. doi:10.1111/imm.12910

8. Beyer M, Mallmann MR, Xue J, et al. High-Resolution Transcriptome of Human Macrophages. PLOS ONE. 2012;7(9):e45466. doi:10.1371/journal.pone.0045466

9. Xue J, Schmidt SV, Sander J, et al. Transcriptome-Based Network Analysis Reveals a Spectrum Model of Human Macrophage Activation. Immunity. 2014;40(2):274–288. doi:10.1016/j.immuni.2014.01.006

10. Spiller KL, Wrona EA, Romero-Torres S, et al. Differential gene expression in human, murine, and cell line-derived macrophages upon polarization. Exp Cell Res. 2016;347(1):1–13. doi:10.1016/j.yexcr.2015.10.017

11. Gerrick KY, Gerrick ER, Gupta A, Wheelan SJ, Yegnasubramanian S, Jaffee EM. Transcriptional profiling identifies novel regulators of macrophage polarization. PLOS ONE. 2018;13(12):e0208602. doi:10.1371/journal.pone.0208602

12. Denisenko E, Guler R, Mhlanga MM, Suzuki H, Brombacher F, Schmeier S. Genome-wide profiling of transcribed enhancers during macrophage activation. Epigenetics Chromatin. 2017;10(1):50. doi:10.1186/s13072-017-0158-9

13. Huang C, Lewis C, Borg NA, et al. Proteomic Identification of Interferon-Induced Proteins with Tetratricopeptide Repeats as Markers of M1 Macrophage Polarization. J Proteome Res. 2018;17(4):1485–1499. doi:10.1021/acs.jproteome.7b00828

14. Li P, Hao Z, Wu J, et al. Comparative Proteomic Analysis of Polarized Human THP-1 and Mouse RAW264.7 Macrophages. Front Immunol. 2021;12:700009. doi:10.3389/fimmu.2021.700009

15. Liu SX, Gustafson HH, Jackson DL, Pun SH, Trapnell C. Trajectory analysis quantifies transcriptional plasticity during macrophage polarization. Sci Rep. 2020;10(1):12273. doi:10.1038/s41598-020-68766-w

16. Muñoz-Rojas AR, Kelsey I, Pappalardo JL, Chen M, Miller-Jensen K. Co-stimulation with opposing macrophage polarization cues leads to orthogonal secretion programs in individual cells. Nat Commun. 2021;12(1):301. doi:10.1038/s41467-020-20540-2

17. Rivera A, Siracusa MC, Yap GS, Gause WC. Innate cell communication kick-starts pathogen-specific immunity. Nat Immunol. 2016;17(4):356–363. doi:10.1038/ni.3375

18. Luecke S, Sheu KM, Hoffmann A. Stimulus-specific responses in innate immunity: Multilayered regulatory circuits. Immunity. 2021;54(9):1915–1932. doi:10.1016/j.immuni.2021.08.018

19. Sheu K, Hoffmann A. in press. Annu Rev Immunol. 2022.

20. Werner SL, Barken D, Hoffmann A. Stimulus Specificity of Gene Expression Programs Determined by Temporal Control of IKK Activity. Science. 2005;309(5742):1857–1861. doi:10.1126/science.1113319

21. Covert MW, Leung TH, Gaston JE, Baltimore D. Achieving Stability of Lipopolysaccharide-Induced NF-κB Activation. Science. 2005;309(5742):1854–1857. doi:10.1126/science.1112304

22. Ashall L, Horton CA, Nelson DE, et al. Pulsatile Stimulation Determines Timing and Specificity of NF-κB-Dependent Transcription. Science. 2009;324(5924):242–246. doi:10.1126/science.1164860

23. Nelson DE, Ihekwaba AEC, Elliott M, et al. Oscillations in NF-κB Signaling Control the Dynamics of Gene Expression. Science. 2004;306(5696):704–708. doi:10.1126/science.1099962

24. Behar M, Hoffmann A. Understanding the temporal codes of intra-cellular signals. Curr Opin Genet Dev. 2010;20(6):684–693. doi:10.1016/j.gde.2010.09.007

25. Sen S, Cheng Z, Sheu KM, Chen YH, Hoffmann A. Gene Regulatory Strategies that Decode the Duration of NFκB Dynamics Contribute to LPS-versus TNF-Specific Gene Expression. Cell Syst. 2020;10(2):169-182.e5. doi:10.1016/j.cels.2019.12.004

26. Tay S, Hughey JJ, Lee TK, Lipniacki T, Quake SR, Covert MW. Single-cell NF-κB dynamics reveal digital activation and analogue information processing. Nature. 2010;466(7303):267–271. doi:10.1038/nature09145

27. Hoffmann A, Levchenko A, Scott ML, Baltimore D. The IκB-NF-κB Signaling Module: Temporal Control and Selective Gene Activation. Science. 2002;298(5596):1241–1245. doi:10.1126/science.1071914

28. Hoffmann A. Immune Response Signaling: Combinatorial and Dynamic Control. Trends Immunol. 2016;37(9):570–572. doi:10.1016/j.it.2016.07.003

29. Cheng QJ, Ohta S, Sheu KM, et al. NF-κB dynamics determine the stimulus specificity of epigenomic reprogramming in macrophages. Science. 2021;372(6548):1349–1353. doi:10.1126/science.abc0269

30. Adelaja A, Taylor B, Sheu KM, Liu Y, Luecke S, Hoffmann A. Six distinct NFκB signaling codons convey discrete information to distinguish stimuli and enable appropriate macrophage responses. Immunity. 2021;54(5):916-930.e7. doi:10.1016/j.immuni.2021.04.011

31. Yang CH, Murti A, Pfeffer SR, Basu L, Kim JG, Pfeffer LM. IFNα/β promotes cell survival by activating NF-κB. Proc Natl Acad Sci. 2000;97(25):13631–13636. doi:10.1073/pnas.250477397

32. Mitchell S, Mercado EL, Adelaja A, et al. An NFκB Activity Calculator to Delineate Signaling Crosstalk: Type I and II Interferons Enhance NFκB via Distinct Mechanisms. Front Immunol. 2019;10:1425. doi:10.3389/fimmu.2019.01425

33. Xu J, Zhou L, Ji L, et al. The REGγ-proteasome forms a regulatory circuit with IκBε and NFκB in experimental colitis. Nat Commun. 2016;7(1):10761. doi:10.1038/ncomms10761

34. Porta C, Rimoldi M, Raes G, et al. Tolerance and M2 (alternative) macrophage polarization are related processes orchestrated by p50 nuclear factor κB. Proc Natl Acad Sci. Published online August 13, 2009. doi:10.1073/pnas.0809784106

35. O’Neill LA, Sheedy FJ, McCoy CE. MicroRNAs: the fine-tuners of Toll-like receptor signalling. Nat Rev Immunol. 2011;11(3):163–175. doi:10.1038/nri2957

36. Quinn SR, O’Neill LA. A trio of microRNAs that control Toll-like receptor signalling. Int Immunol. 2011;23(7):421–425. doi:10.1093/intimm/dxr034

37. Curtale G, Mirolo M, Renzi TA, Rossato M, Bazzoni F, Locati M. Negative regulation of Toll-like receptor 4 signaling by IL-10–dependent microRNA-146b. Proc Natl Acad Sci. 2013;110(28):11499–11504. doi:10.1073/pnas.1219852110

38. Tang B, Xiao B, Liu Z, et al. Identification of MyD88 as a novel target of miR-155, involved in negative regulation of Helicobacter pylori-induced inflammation. FEBS Lett. 2010;584(8):1481–1486. doi:10.1016/j.febslet.2010.02.063

39. O’Connell RM, Taganov KD, Boldin MP, Cheng G, Baltimore D. MicroRNA-155 is induced during the macrophage inflammatory response. Proc Natl Acad Sci. 2007;104(5):1604–1609. doi:10.1073/pnas.0610731104

40. Jablonski KA, Gaudet AD, Amici SA, Popovich PG, Guerau-de-Arellano M. Control of the Inflammatory Macrophage Transcriptional Signature by miR-155. PLOS ONE. 2016;11(7):e0159724. doi:10.1371/journal.pone.0159724

41. Taganov KD, Boldin MP, Chang KJ, Baltimore D. NF-κB-dependent induction of microRNA miR-146, an inhibitor targeted to signaling proteins of innate immune responses. Proc Natl Acad Sci. 2006;103(33):12481–12486. doi:10.1073/pnas.0605298103

42. Testa U, Pelosi E, Castelli G, Labbaye C. miR-146 and miR-155: Two Key Modulators of Immune Response and Tumor Development. Non-Coding RNA. 2017;3(3):22. doi:10.3390/ncrna3030022

43. Ruedl C, Khameneh HJ, Karjalainen K. Manipulation of immune system via immortal bone marrow stem cells. Int Immunol. 2008;20(9):1211–1218. doi:10.1093/intimm/dxn079

44. Sakoe H, Chiba S. Dynamic programming algorithm optimization for spoken word recognition. IEEE Trans Acoust Speech Signal Process. 1978;26(1):43–49. doi:10.1109/TASSP.1978.1163055

45. Cuturi M, Blondel M. Soft-DTW: a differentiable loss function for time-series. In: Proceedings of the 34th International Conference on Machine Learning - Volume 70. ICML’17. JMLR.org; 2017:894–903.

46. Tavenard R, Faouzi J, Vandewiele G, et al. Tslearn, A Machine Learning Toolkit for Time Series Data. J Mach Learn Res. 2020;21(118):1–6.

47. Lane K, Andres-Terre M, Kudo T, Monack DM, Covert MW. Escalating Threat Levels of Bacterial Infection Can Be Discriminated by Distinct MAPK and NF-κB Signaling Dynamics in Single Host Cells. Cell Syst. 2019;8(3):183-196.e4. doi:10.1016/j.cels.2019.02.008

48. Hochreiter S, Schmidhuber J. Long Short-Term Memory. Neural Comput. 1997;9(8):1735–1780. doi:10.1162/neco.1997.9.8.1735

49. Van Houdt G, Mosquera C, Nápoles G. A review on the long short-term memory model. Artif Intell Rev. 2020;53(8):5929–5955. doi:10.1007/s10462-020-09838-1

50. Ramsay JO, Silverman BW. Functional Data Analysis. 2nd ed. Springer; 2005.

51. Sheu KM, Luecke S, Hoffmann A. Stimulus-specificity in the responses of immune sentinel cells. Curr Opin Syst Biol. 2019;18:53–61. doi:10.1016/j.coisb.2019.10.011

52. Lawrence T, Natoli G. Transcriptional regulation of macrophage polarization: enabling diversity with identity. Nat Rev Immunol. 2011;11(11):750–761. doi:10.1038/nri3088

53. Takeuch O, Akira S. Epigenetic control of macrophage polarization. Eur J Immunol. 2011;41(9):2490–2493. doi:10.1002/eji.201141792

54. Ivashkiv LB. Epigenetic regulation of macrophage polarization and function. Trends Immunol. 2013;34(5):216–223. doi:10.1016/j.it.2012.11.001

55. Gosselin D, Glass CK. Epigenomics of macrophages. Immunol Rev. 2014;262(1):96–112. doi:10.1111/imr.12213

56. Lavin Y, Winter D, Blecher-Gonen R, et al. Tissue-resident macrophage enhancer landscapes are shaped by the local microenvironment. Cell. 2014;159(6):1312–1326. doi:10.1016/j.cell.2014.11.018

57. Van den Bossche J, Neele AE, Hoeksema MA, de Winther MPJ. Macrophage polarization: the epigenetic point of view. Curr Opin Lipidol. 2014;25(5):367–373. doi:10.1097/MOL.0000000000000109

58. Chen S, Yang J, Wei Y, Wei X. Epigenetic regulation of macrophages: from homeostasis maintenance to host defense. Cell Mol Immunol. 2020;17(1):36–49. doi:10.1038/s41423-019-0315-0

59. Cochain C, Vafadarnejad E, Arampatzi P, et al. Single-Cell RNA-Seq Reveals the Transcriptional Landscape and Heterogeneity of Aortic Macrophages in Murine Atherosclerosis. Circ Res. 2018;122(12):1661–1674. doi:10.1161/CIRCRESAHA.117.312509

60. MacParland SA, Liu JC, Ma XZ, et al. Single cell RNA sequencing of human liver reveals distinct intrahepatic macrophage populations. Nat Commun. 2018;9:4383. doi:10.1038/s41467-018-06318-7

61. Seidman JS, Troutman TD, Sakai M, et al. Niche-Specific Reprogramming of Epigenetic Landscapes Drives Myeloid Cell Diversity in Nonalcoholic Steatohepatitis. Immunity. 2020;52(6):1057-1074.e7. doi:10.1016/j.immuni.2020.04.001

62. Liao M, Liu Y, Yuan J, et al. Single-cell landscape of bronchoalveolar immune cells in patients with COVID-19. Nat Med. 2020;26(6):842–844. doi:10.1038/s41591-020-0901-9

63. Zhang Q, Cheng S, Wang Y, et al. Interrogation of the microenvironmental landscape in spinal ependymomas reveals dual functions of tumor-associated macrophages. Nat Commun. 2021;12(1):6867. doi:10.1038/s41467-021-27018-9

64. Mei Y, Xiao W, Hu H, et al. Single-cell analyses reveal suppressive tumor microenvironment of human colorectal cancer. Clin Transl Med. 2021;11(6):e422. doi:10.1002/ctm2.422

65. Kinnunen PC, Luker KE, Luker GD, Linderman JJ. Computational methods for characterizing and learning from heterogeneous cell-signaling data. Curr Opin Syst Biol. Published online May 4, 2021. doi:10.1016/j.coisb.2021.04.009

66. Singh A, Marcoline FV, Veshaguri S, et al. Protons in small spaces: Discrete simulations of vesicle acidification. PLOS Comput Biol. 2019;15(12):e1007539. doi:10.1371/journal.pcbi.1007539

67. Alberto Mantovani, Antonio Sica, Locati M. Macrophage Polarization Comes of Age. Immunity. 2005;23(4):344–346. doi:10.1016/j.immuni.2005.10.001

68. Sica A, Mantovani A. Macrophage plasticity and polarization: in vivo veritas. J Clin Invest. 2012;122(3):787–795. doi:10.1172/JCI59643

69. Mosser DM, Edwards JP. Exploring the full spectrum of macrophage activation. Nat Rev Immunol. 2008;8(12):958–969. doi:10.1038/nri2448

70. Chinetti-Gbaguidi G, Colin S, Staels B. Macrophage subsets in atherosclerosis. Nat Rev Cardiol. 2015;12(1):10–17. doi:10.1038/nrcardio.2014.173

71. Nakai K. Multiple roles of macrophage in skin. J Dermatol Sci. 2021;104(1):2–10. doi:10.1016/j.jdermsci.2021.08.008

72. Guillemin A, Stumpf MPH, Guillemin A, Stumpf MPH. Non-equilibrium statistical physics, transitory epigenetic landscapes, and cell fate decision dynamics. Math Biosci Eng. 2020;17(6):7916–7930. doi:10.3934/mbe.2020402

73. Martin M. Cutadapt removes adapter sequences from high-throughput sequencing reads. EMBnet.journal. 2011;17(1):10–12. doi:10.14806/ej.17.1.200

74. Schmieder R, Edwards R. Quality control and preprocessing of metagenomic datasets. Bioinformatics. 2011;27(6):863–864. doi:10.1093/bioinformatics/btr026

75. Dobin A, Davis CA, Schlesinger F, et al. STAR: ultrafast universal RNA-seq aligner. Bioinformatics. 2013;29(1):15–21. doi:10.1093/bioinformatics/bts635

76. Danecek P, Bonfield JK, Liddle J, et al. Twelve years of SAMtools and BCFtools. GigaScience. 2021;10(2):giab008. doi:10.1093/gigascience/giab008

77. Liao Y, Smyth GK, Shi W. featureCounts: an efficient general purpose program for assigning sequence reads to genomic features. Bioinformatics. 2014;30(7):923–930. doi:10.1093/bioinformatics/btt656

78. Freedman D, Diaconis P. On the histogram as a density estimator:L2 theory. Z Für Wahrscheinlichkeitstheorie Verwandte Geb. 1981;57(4):453–476. doi:10.1007/BF01025868

79. Pedregosa F, Varoquaux G, Gramfort A, et al. Scikit-learn: Machine Learning in Python. J Mach Learn Res. 2011;12(85):2825–2830.

80. Abadi M, Barham P, Chen J, et al. TensorFlow: A System for Large-Scale Machine Learning. In: ; 2016:265–283. Accessed November 12, 2021. https://www.usenix.org/conference/osdi16/technical-sessions/presentation/abadi

81. Chollet F. Keras.; 2015. Accessed November 12, 2021. https://keras.io

82. Suárez A, Torrecilla JL, Berrocal MC, et al. Scikit-Fda. Grupo de Aprendizaje Automático - Universidad Autónoma de Madrid; 2019.

83. McInnes L, Healy J, Melville J. UMAP: Uniform Manifold Approximation and Projection for Dimension Reduction. ArXiv180203426 Cs Stat. Published online September 17, 2020. Accessed February 24, 2022. http://arxiv.org/abs/1802.03426

